# How genotype-by-environment interactions can maintain variation in mutualisms

**DOI:** 10.1101/2024.07.19.604331

**Authors:** Christopher I. Carlson, Megan E. Frederickson, Matthew M. Osmond

## Abstract

Coevolution requires reciprocal genotype-by-genotype (GXG) interactions for fitness, which occur when the fitnesses of interacting species depend on the match between their genotypes. However, in mutualisms, when GXG interactions are mutually beneficial, simple models predict that positive feedbacks will erode genetic variation, weakening or eliminating the GXG interactions that fuel ongoing coevolution. This is inconsistent with the ample trait and fitness variation observed within real-world mutualisms. Here, we explore how genotype-by-environment (GXE) interactions, which occur when different genotypes respond differently to different environments, maintain variation in mutualisms. We employ a game theoretic model in which the fitnesses of two partners depend on mutually beneficial GXG and GXE interactions. Variation is maintained via migration-selection balance when GXE interactions are slightly stronger than GXG interactions or when they are much stronger than GXG interactions for just one partner. However, unexpectedly, when GXE interactions are much stronger than GXG interactions for both partners and dispersal is high, genotypically mismatched partners can fix, eroding variation and leading to apparent maladaptation between partners. We parameterize our model using data from three published reciprocal transplant experiments and find that the observed strengths of GXE interactions can maintain or erode variation in mutualisms via these mechanisms.

## Introduction

The world is replete with biological variation that appears to have arisen via coevolution. However, in mutualisms, simple models suggest that coevolution will erode rather than maintain or promote variation. Coevolution occurs when the fitnesses of two or more interacting species depend on the match between their genotypes via genotype-by-genotype, or GXG, interactions. Mutually beneficial GXG interactions occur when a match between partner genotypes enhances the fitness of both partners, for example, as when rhizobia evolve to benefit their local legume genotype (Batstone et al., 2020). Mutually beneficial GXG interactions occur in many mutualisms, including between legumes and rhizobia (Batstone et al., 2022, 2020; Heath et al., 2012), plants and microbiomes (O’Brien et al., 2022), plants and mycorrhizal fungi (Hoeksema, 2010), plants and pollinators (Pauw et al., 2009; Peralta et al., 2020), and corals and algae (Parkinson and Baums, 2014). However, in mutualisms, when GXG interactions are mutually beneficial, simple coevolutionary models predict that positive frequency dependent natural selection will erode genetic variation (Carlson et al., 2022; Lively and Wade, 2022; Nuismer, 2017). Given widespread genetic variation and GXG interactions in mutualisms (Bayliss et al., 2019; Friesen, 2012; Harrison et al., 2017; Hoeksema, 2010; Pellmyr and Huth, 1994; Poulsen et al., 2009), understanding the processes that generate the variation fueling ongoing coevolution is a longstanding challenge for mutualism theory (Heath and Stinchcombe, 2014; Hoeksema, 2010; Stoy et al., 2020; Thompson, 1999).

One possible explanation for how genetic variation is maintained in mutualisms is that the same coevolutionary processes that maintain variation in antagonistic interactions also maintain variation in mutualisms. For example, selection to cheat, i.e., to improve the relative fitness of a focal genotype at the expense of the relative fitness of its partner (Jones et al., 2015), could lead to conflicts of interest between mutualists (Ferriere et al., 2002; Kiers et al., 2003; Wechsler and Bascompte, 2022). Significant theoretical and experimental attention has been devoted to understanding how this conflict promotes variation in mutualism (Carlson et al., 2022; Gano-Cohen et al., 2019; Gomulkiewicz et al., 2003; Jones et al., 2015); however, there is strong empirical evidence that many mutualists have aligned fitnesses, meaning selection favors cooperators, not cheaters (Frederickson, 2017; Friesen, 2012; Jones et al., 2015). Therefore, understanding how genetic variation is maintained in mutualisms when GXG interactions are mutually beneficial is crucial (Frederickson, 2017, 2020; Friesen, 2012). Another factor that may maintain genetic variation in mutualism is genotype-by-environment (GXE) interactions, which occur when different genotypes respond differently to different environments (Heath and Stinchcombe, 2014; Vaidya and Stinchcombe, 2020). Here, we explore mathematically how GXE interactions, which can lead to variation in any trait, can promote variation in a mutualism in which partners have aligned fitness interests.

Empirical research has found that GXE interactions occur in many, and probably most, mutualisms. For example, GXE interactions mediate the fitness benefits derived from mutualism in plant-microbe interactions (Petipas et al., 2020; Vaidya and Stinchcombe, 2020), plant mycorrhizal symbiosis (Piculell et al., 2008), and pollination mutualisms (Thompson and Pellmyr, 1992). GXE interactions occur because of changes in the degree to which different genotypes exhibit phenotypic plasticity between environments (Scheiner, 1993). These interactions are thought to maintain genetic variation between environments when plasticity leads the ‘best’ genotype to change, resulting in changes in the rank order of genotype fitnesses (Johnson, 2007; Vaidya and Stinchcombe, 2020). Species distributed across a heterogeneous landscape then experience balancing selection mediated by GXE interactions that promotes genetic variation between environments (Gillespie and Turelli, 1989; Heath and Nuismer, 2014; Thompson, 2005) because there is variable selection favoring genotypes that are the fittest in their preferred localities. Furthermore, when there is dispersal between environments, GXE can also promote variation within patches. For example, a genotype that is adapted to one patch could be stably maintained via dispersal in a patch to which it is maladapted, reflecting a migration-selection balance (reviewed in Felsenstein, 1976).

Frequency dependence, the relative strengths of GXE vs. GXG interactions, and asymmetric dispersal complicate predictions of the role of GXE interactions in promoting genetic variation for mutualism. In one-species models, GXE maintains variation in an intuitive fashion as genotypes experience consistent changes in fitness between environments (Gillespie and Turelli, 1989). But in mutualisms, GXG interactions lead to positive frequency dependent selection that could potentially overwhelm GXE interactions for fitness between environments. Indeed, a two patch population genetic model of mutualism with mutually beneficial GXG interactions (Nuismer, 2017, chapter 8) suggests GXE interactions are not strong enough to maintain variation in the face of positive frequency dependent selection. However, this model incorporates several assumptions that make GXE interactions unlikely to maintain variation in the first place. The analysis of the model assumes that migration between different environments is frequent and that GXG interactions for fitness are stronger than GXE interactions for both partners. Real mutualisms may involve a variety of relative strengths of GXG vs. GXE interactions that may influence evolutionary dynamics. The strengths of these interactions could even be asymmetric between partners. For example, when a GXE interaction is strong in one partner, but a GXG interaction is strong in the other, it is unclear whether subsequent allele frequency dynamics will reflect selection to be locally adapted to the environment, or selection to match a partner’s genotype, or some balance of the two. Finally, the role of GXE interactions in maintaining variation in mutualism is complicated because mutualistic partners often have different dispersal rates; e.g., beneficial microbes could have dispersal rates far above or below that of their plant hosts (Baltrus, 2020; Custer et al., 2022). It is unclear how the eco-evolutionary dynamics of mutualism depend on these two dispersal rates; they may be best explained by the smaller dispersal rate, the larger rate, or by the mean (Carlson et al., 2022). Given these complexities, here we build a mathematical model to help us understand the role of GXE interactions in maintaining variation in mutualisms.

Our model explores to what extent GXE interactions promote genetic variation in mutualism when mutualistic partners exhibit mutually beneficial GXG interactions and disperse between environments. In the model, the two genotypes within one species interact with the two genotypes in the other species in each of the two distinct environments in proportion to their frequencies. The benefit derived by mutualists from mutualism depends on the match between partner genotypes (GXG interaction) and the match between a given individual’s genotype and its environment (GXE interaction). We find that GXE interactions maintain significant variation in mutualism when intermediate in strength in one or both partners, when a GXE interaction is strong in just one partner, or when one partner disperses at a low rate. Strong GXE or GXG interactions and high dispersal in both partners can erode variation and promote apparent maladaptation to local environments or mutualistic partners. These results suggest that GXE interactions can maintain variation in mutualisms but primarily do so when they are of similar magnitude to GXG interactions, or when there is asymmetry in their strength or in dispersal across partners. Finally, we parameterized our model’s payoffs using data from reciprocal transplant studies of plant-microbe, plant-mycorrhizae, and *Drosophila melanogaster*-microbiome interactions and find relative strengths of GXE interactions that are predicted to maintain variation in the wild.

## Methods

Our model uses a discrete-strategy game-theoretic framework with two species that we refer to as hosts and symbionts. This general framework can encompass many horizontally transmitted mutualisms ranging from the symbiosis between plants and mycorrhizal fungi, to diffuse partnerships such as seed dispersal or cleaner fish-client fish mutualisms (Bshary and Bronstein, 2011). In the model, there are two host genotypes, host 1 and host 2, and two symbiont genotypes, symbiont 1 and symbiont 2. These species form an obligate mutualistic partnership while dispersing between two environments, environment 1 and environment 2. Hosts and symbionts engage in an evolutionary game, where partners exchange rewards that increase their fitness payoffs. These payoffs depend on: 1) the match between the host and symbiont genotypes, 2) the match between the host and its environment, and 3) the match between the symbiont and its environment. Each genotype receives one of four total payoffs, with *H* or *S* denoting whether the payoff is received by a host or symbiont, respectively:

*H_g_*/*S_g_* : Only host and symbiont *genotypes* match (host- and symbiont-environment mismatch)

*H_h_*/*S_h_* : Only the *host* matches its environment (symbiont-environment & genotypic mismatch)

*H_s_*/*S_s_* : Only the *symbiont* matches its environment (host-environment & genotypic mismatch)

*H_m_*/*S_m_* : There is a genotypic, host-environment, & symbiont-environment *match*

Note that, for simplicity, *H_i_* and *S_i_*are symmetrical with respect to genotype (e.g., *H_m_* for host 1 is the same as *H_m_* for host 2).

Next, we derive expressions for the average payoff (i.e., fitness) of each genotype in each environment. We assume that there is no partner choice; hosts and symbionts interact according to their frequency in each environment, and individuals cannot discriminate between partners based on their quality. Our model reflects interactions among many different partner genotypes (e.g. plantpollinator or seed dispersal mutualisms) or partnerships where each host has a single symbiont that it acquires from a well mixed local community (e.g., coral-algal symbiosis). The model applies to both kinds of mutualisms because the fitness of each focal genotype is equal to the frequency of its two partners weighted by the fitness payoff (see section S.1).

It is enough to track the frequency of genotype 1 in each species and environment since there are only two genotypes whose frequencies sum to 1. We denote these frequencies *h_i_* and *s_i_* for the host or symbiont, respectively, in environment *i* (*i* = 1 or *i* = 2). Let *w_ji_*(*s_i_*) and *v_ji_*(*h_i_*) denote the fitnesses of genotype *j* in environment *i* for host and symbionts, respectively. For example, in environment 1, host 1 receives *H_m_* when it interacts with symbiont 1 and *H_h_* when it interacts with symbiont 2, giving *w*_11_(*s*_1_) = *s*_1_ *H_m_*+ (1 − *s*_1_)*H_h_*. For all expressions, see section S.1.

Let the fitness advantage of genotype 1 in environment *i* be Δ*w_i_*(*s_i_*) = *w*_1*i*_(*s_i_*) − *w*_2*i*_(*s_i_*) and Δ*v_i_*(*h_i_*) = *v*_1*i*_(*h_i_*) − *v*_2*i*_(*h_i_*) for hosts and symbionts, respectively. It turns out that we can express these fitness advantages in terms of differences between payoffs and therefore that these payoff differences dictate the model’s dynamics. Specifically, we define four payoff differences:

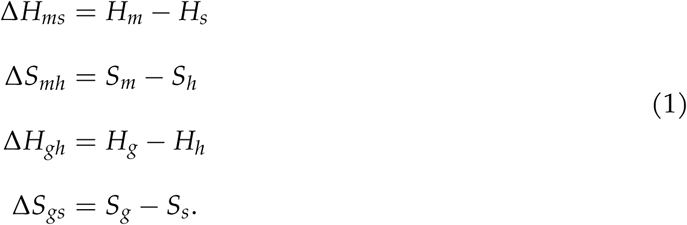

The first two differences represent the difference between a perfect match and interacting with a partner that matches its environment. The latter two differences represent the difference between matching a partner’s genotype and matching the environment. The fitness advantages for genotype 1 hosts and symbionts across environments 1 and 2 are then

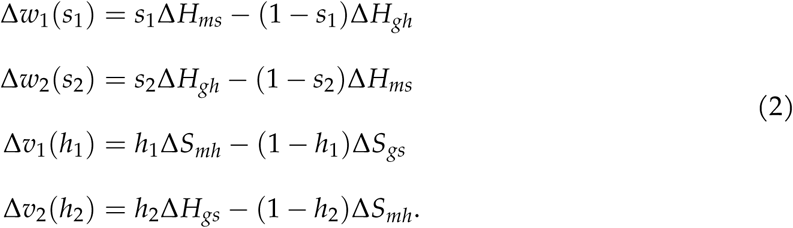

With these fitness advantages in hand, we can write dynamical expressions for the change in the frequency of each host and each symbiont within each environment using the replicator equation (Taylor and Jonker, 1978). In these expressions, a genotype increases in frequency in proportion to its fitness advantage. We also let hosts and symbionts disperse between environments in direct proportion to their frequencies. For simplicity we start by assuming that both species disperse at rate *d*, but relax this later. Throughout, we assume the dispersal rate from environment 1 to 2 is the same as the rate from environment 2 to 1, e.g., symmetric dispersal between patches of equal size.

Our expressions for the rate of change in frequency of each genotype in each environment are thus

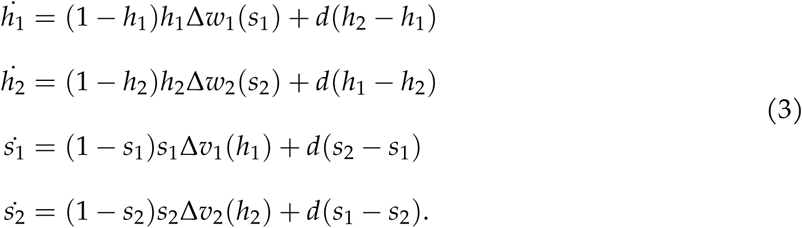

We first find all boundary equilibria, where genotypes either fix or go extinct (i.e., the frequency of each genotype is 0 or 1), and evaluate their stability. Given the non-linearities in equation 3, it is impossible to analytically solve for all internal equilibria with nonzero *d*, so we instead approximate these equilibria and their stability assuming weak dispersal. We then compare these approximations to full numerical simulations. Next, we consider a scenario in which hosts and symbionts disperse at different rates, *d_h_* and *d_s_*, respectively, and analyze the model similarly. See our online supplemental materials for details of the mathematical analysis and https://github.com/chrisiancarlson/GXE-mutualism for code and simulation details. Finally, we parameterize the model’s payoffs using data from plant-arbuscular mycorrhizal (AM) fungi, plant-microbe, and fly-microbiome interactions to demonstrate that our approach can be used to examine how GXE interactions are expected to maintain variation in real-world mutualisms.

## Results

### Biologically relevant equilibria

The model always features two sets of biologically relevant boundary equilibria that can be solved exactly. We refer to the first set of boundary equilibria as a *global genotypic match* where one host and its preferred symbiont fix in both environments (i.e., host 1 fixes everywhere with symbiont 1 or host 2 fixes everywhere with symbiont 2; see section S.2.1 for more details). The second set of boundary equilibria is a *global genotypic mismatch* where a host and its mismatched symbiont fix in both environments (i.e., host 1 fixes everywhere with symbiont 2 or host 2 fixes everywhere with symbiont 1; see section S.2.2).

There are also two biologically relevant internal equilibria when dispersal is small. The first of these is a *migration-selection balance* equilibrium, where genotypically-matched hosts and symbionts are frequent in their preferred environment and are maintained at a low frequency in their mismatched environment by dispersal (see section S.2.3). The second equilibrium is a *reverse migration-selection balance* in which a genotypically matched host and symbiont pair is frequent in its mismatched environment, but infrequent in its matched environment (see section S.2.4). As dispersal goes to 0, one pair of genotypically matched hosts and symbionts fix in environment 1, and the other pair of genotypically matched hosts and symbionts fix in environment 2.

### Stability

We next examine the local stability of these equilibria. Assuming that a full match (i.e., the partners match each other and their environment) gives a greater payoff than having only a partner that matches the environment, Δ*H_ms_ >* 0 and Δ*S_mh_ >* 0, we find that there are four different cases depending on the sign and magnitude of Δ*H_gh_*and Δ*S_gs_*(see sections S.2.1-S.2.4 for mathematical details). These four cases can be understood in terms of the reaction norms of a focal genotype between partner genotypes and between environments or, equivalently, the strength of GXE interactions relative to GXG interactions.

In the first case, GXE interactions are weak enough such that both Δ*H_gh_*and Δ*S_gs_*are positive. This implies that the reaction norms comparing fitness between genotype pairings have slopes of opposite sign and cross, which changes the rank order fitness between partner genotypes (figure 1b, strong GXG). Meanwhile the reaction norms between environments can be parallel (no GXE) or have slopes of the same sign and not cross (figure 1a, weak GXE).

**Figure 1:**
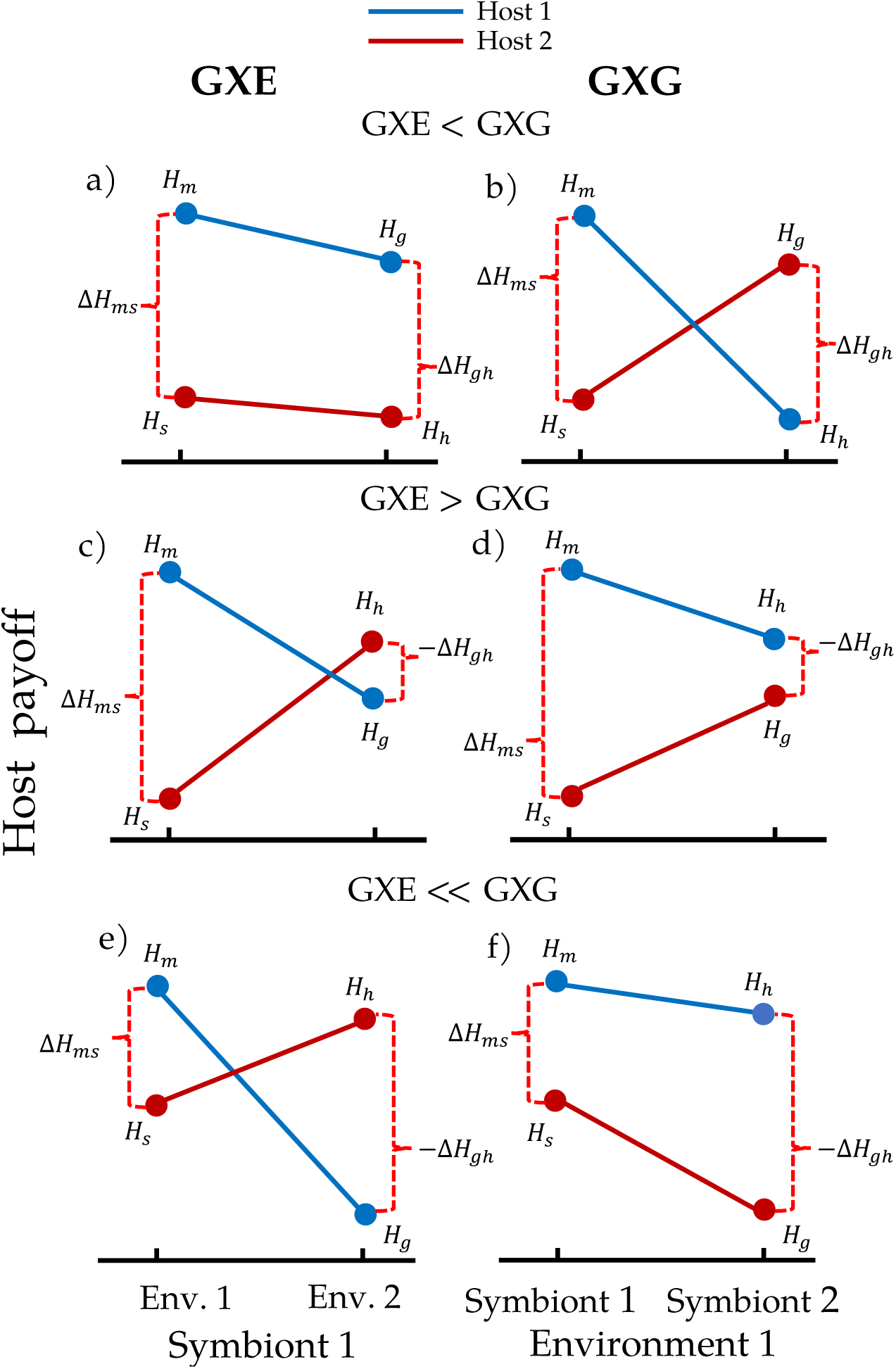
The relationship between fitness payoffs and the relative strength of GXE vs. GXG interactions. This figure depicts each host genotype’s reaction norm as a function of environment when partnered with symbiont 1 (panels a, c and e) and as as a function of symbiont type in environment 1 (panels b, d, and f). Line colors correspond to host genotype. The rows are the relative strengths of GXE vs. GXG interactions, with GXE < GXG in the top row (a and b), GXE > GXG in the middle row (c and d), and GXE *>>* GXG in the bottom row (e and f). The red, dashed brackets denote the value and sign of the payoff advantage terms (Δ*H_ms_* or Δ*H_gh_*).

In the second case, GXE interactions are strong enough to cause at least one of Δ*H_gh_* or Δ*S_gs_* to become negative but weak enough such that both Δ*H_ms_*+ Δ*H_gh_*and Δ*S_mh_*+ Δ*S_gs_*are still positive. Now the reaction norms between partner genotypes for the species with a negative Δ*H_gh_*or Δ*S_gs_*still have opposite signs but no longer cross (figure 1d, moderate GXG) while their reaction norms between environments have slopes of opposite sign and cross (figure 1c, strong GXE).

In the third case, GXE interactions are again strong enough to cause at least one of Δ*H_gh_* and Δ*S_gs_*to become negative but now also strong enough to make one of Δ*H_ms_*+ Δ*H_gh_*and Δ*S_mh_*+ Δ*S_gs_*negative. Now for one species the slopes of the reaction norms between genotypes have the same signs and do not cross (figure 1f, weak GXG) while the reaction norms between environments still cross (figure 1e, strong GXE).

In the fourth case, GXE interactions are strong enough to cause both Δ*H_gh_*and Δ*S_gs_*to become sufficiently negative that both Δ*H_ms_* + Δ*H_gh_* and Δ*S_mh_* + Δ*H_gs_* are negative. Now both species have reaction norms as just described (figure 1e,f).

We present our analysis of each case in order of increasing strength of GXE relative to GXG interactions, from case one to case four. The results are summarized in Table 1.

**Table 1:**
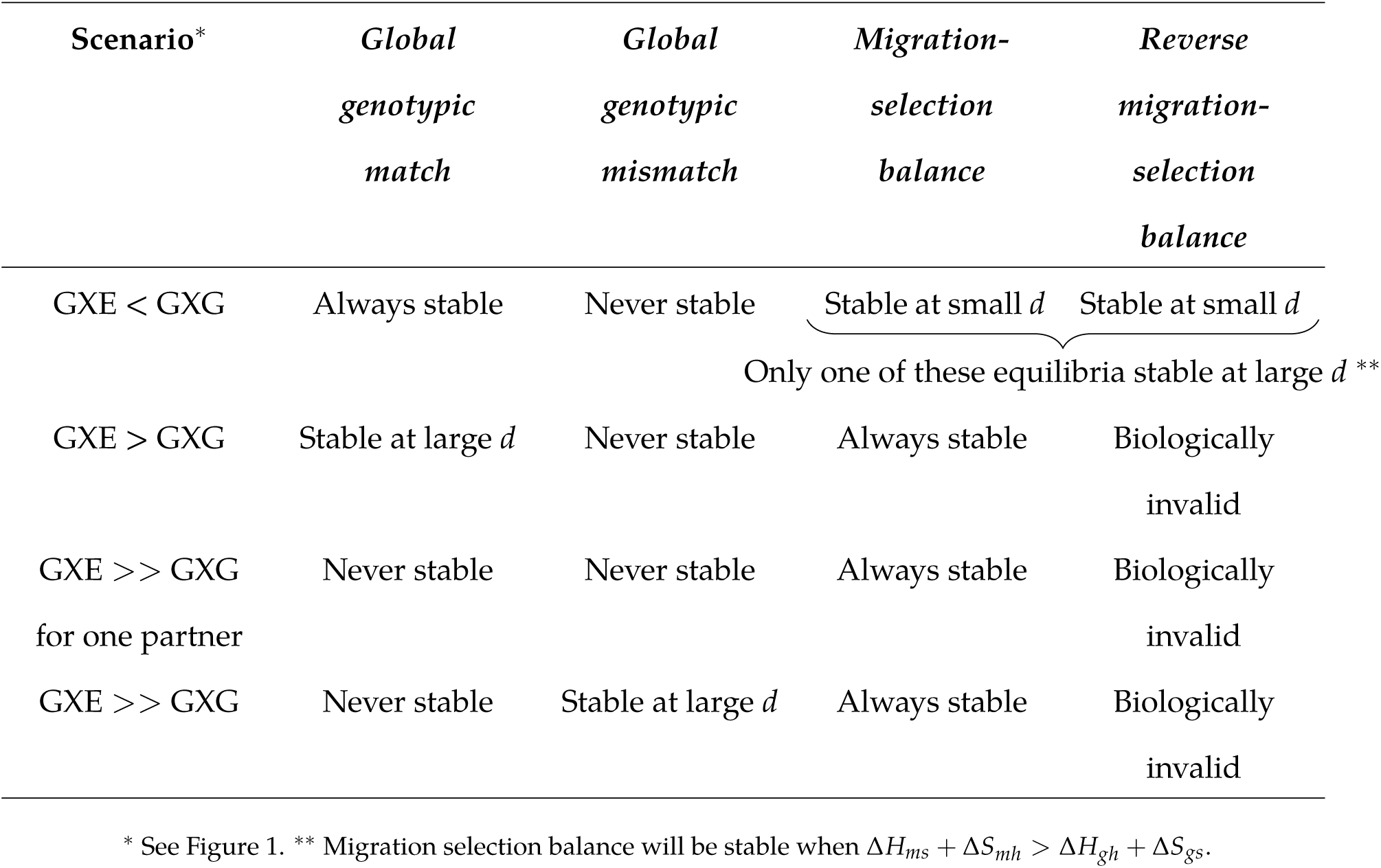
Summary of equilibria and their local stability.

#### When matching genotypes is more advantageous than matching the environment (GXE < GXG)

When GXE interactions are weak (or absent) and GXG interactions are strong (figure 1a-b), the *global genotypic match*, *migration-selection balance*, and *reverse migration-selection balance* equilibria can all be locally stable (figure 2). The latter two of these equilibria are polymorphic, meaning that even when GXE interactions are weak or absent, spatial structure can maintain variation in interacting mutualists. Next, we explain the intuition behind the stability of each of these equilibria.

**Figure 2:**
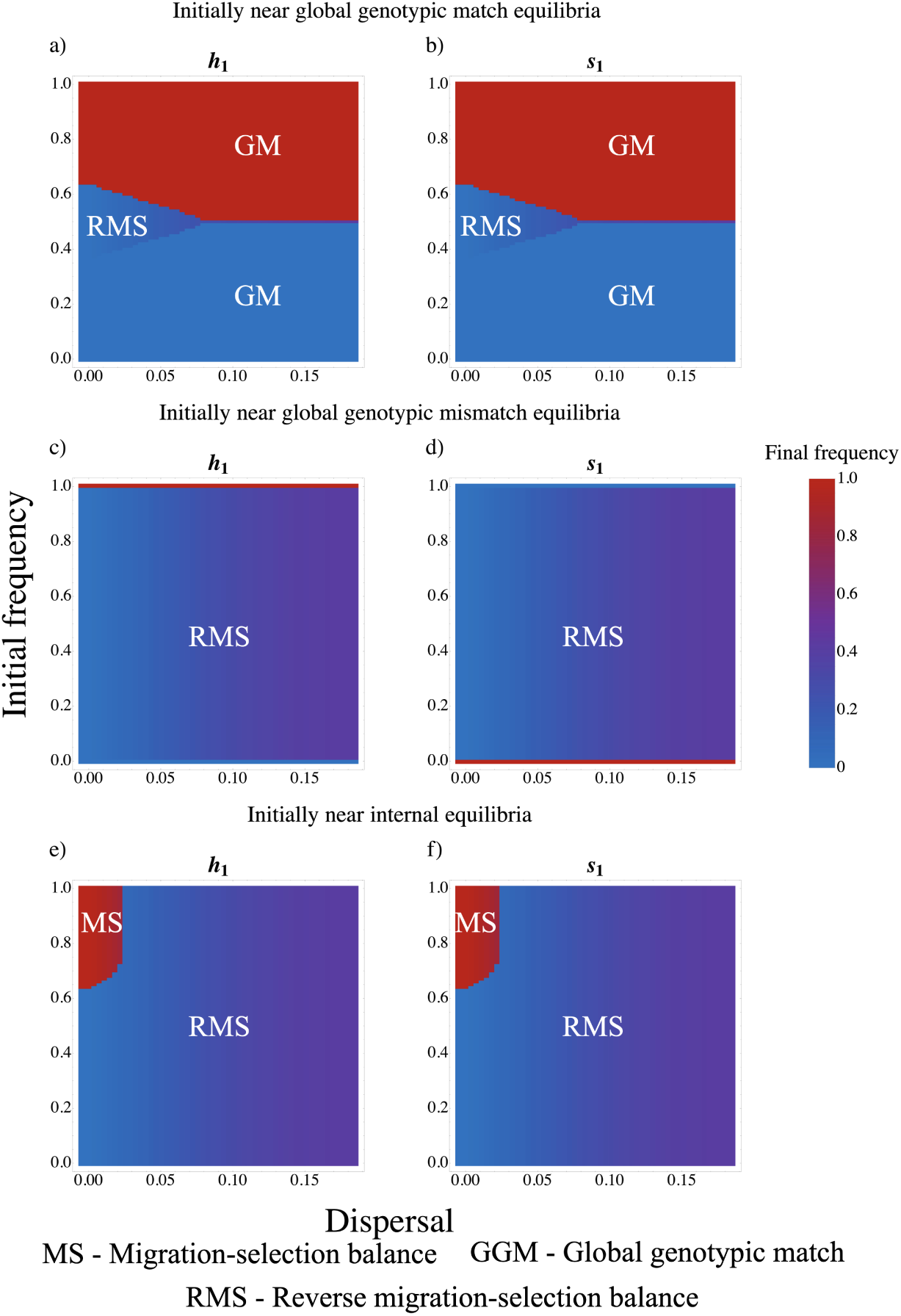
Results of simulations where GXE < GXG. Here Δ*H_ms_* = Δ*S_mh_* = 0.2 and Δ*H_gh_* = Δ*S_gs_* = 0.35. Each panel gives the frequency of genotype 1 in environment 1, *h*_1_ and *s*_1_ for hosts and symbionts, respectively, at *t* = 10, 000. The *x* axis is dispersal rate, *d*, and the *y* axis is the initial frequency of host genotype 1 in environment 1, *h*_1_. The other initial frequencies differ by row. In the first row, the initial frequencies of genotype 1 increase uniformly across both environments and species, *h*_1_ = *h*_2_ = *s*_1_ = *s*_2_. In the second row, as the initial frequencies of genotype 1 in the host increase the initial frequencies in the symbiont decrease, *h*_1_ = *h*_2_ = 1 − *s*_1_ = 1 − *s*_2_. In the third row, as the initial frequencies of genotype 1 in environment 1 increase the initial frequencies in environment 2 decrease, *h*_1_ = *h*_2_ = 1 − *s*_1_ = 1 − *s*_2_. mismatching the environment is still relatively low.

First, the *global genotypic match* equilibria are always locally stable (figure 2a-b) because when GXG interactions are strong, matching a mutualistic partner’s genotype is more beneficial than matching the environment (figure 1a). At these equilibria, GXG interactions completely erode genetic variation in the mutualistic partners both within and between environments, recovering the behavior of a non-spatial, matching allele model of mutualism (e.g., Nuismer, 2017, chapter 2).

Second, the *migration-selection balance* equilibrium is always locally stable under our approximation that dispersal is rare (figure 2e-f). The strong positive frequency dependence imparted by GXG interactions maintains the locally adapted pair within each environment, with fixation prevented by dispersal. This reflects an outcome where, despite weak or even absent GXE interactions, spatial structure maintains both genotypes locally within each environment.

The *reverse migration-selection balance* equilibrium is also always locally stable under our approximation that dispersal is rare (figure 2e-f). Like *migration-selection balance*, this equilibrium is locally stable because there is positive frequency dependence in the provisioning of the high benefit from matching genotypes and because mismatching genotypes causes a change in the rank ordering of payoffs (strong GXG, figure 1a). For example, when a matching host-symbiont pair is frequent in a given environment, the infrequent genotype introduced via dispersal is primarily interacting with its genotypically-mismatched partner. When matching a partner is much more important to fitness than matching the environment, the infrequent genotype cannot achieve high frequency because of the high cost of a genotypic mismatch even if it matches the environment. This equilibrium demonstrates that even if GXE interactions are weak to nonexistent, positive frequency dependence and spatial structure can maintain polymorphism in the form of local maladaptation. Without dispersal all four equilibria are locally stable, driven by positive frequency dependence within environments. Increasing dispersal reduces the range of initial conditions that lead the system toward the internal equilibria (figure 2a-b), thereby decreasing the probability of any variation. Higher dispersal rates increase the shuffling of genotypes across environments and partners, disrupting local adaptation and frequency-dependence and thus favoring whichever pair of genotypes has the highest average fitness across environments. However, if the system is able to reach one of the internal equilibria, increased dispersal increases variation within any one environment by bringing in more of the infrequent genotypes (figure 2a-f).

While our analysis of the internal equilibria assumes weak dispersal, simulations suggest that at higher levels of dispersal one of the internal equilibria maintains stability (figure 2e-f). Which equilibrium remains stable depends on the inequality (Δ*H_ms_* + Δ*S_mh_*) ≶ (Δ*H_gh_* + Δ*S_gs_*). If (Δ*H_ms_* + Δ*S_mh_*) *>* (Δ*H_gh_* + Δ*S_gs_*) *migration-selection balance* is stable at high dispersal because the average fitness advantage from matching a preferred environment is higher than the advantage in the mismatched environment. If (Δ*H_ms_* + Δ*S_mh_*) *<* (Δ*H_gh_* + Δ*S_gs_*) then *reverse migration-selection balance* is stable at high dispersal because now there is a greater average fitness advantage for hosts and symbionts in their mismatched environment.

#### When matching the environment is slightly more advantageous than matching genotypes (GXE > GXG)

When GXE is strong and GXG is moderate (figure 1c-d) for at least one partner, we find that the *migration-selection balance* equilibrium is stable at low and high dispersal, while the *global genotypic match* equilibria are locally stable at high dispersal (figure S.1). The stability of these equilibria indicates that moderately strong GXE interactions promote genetic variation via migration-selection balance, especially when dispersal is rare.

The *migration-selection balance* equilibrium can be locally stable because the local benefits of matching the environment overwhelm the influx of mismatched genotypes from the opposite environment. As in the previous section (GXE<GXG), this equilibrium is always locally stable under our assumption of rare dispersal because there is always an advantage to matching the environment (see figure 1d). Simulations suggest that this local advantage maintains the stablitity of this equilibrium even at high rates of dispersal (figure S1e-f).

The *global genotypic match* equilibria only become stable once dispersal becomes sufficiently high. This is because, for high enough dispersal and the right initial conditions (see figure S1ab), the global advantage of matching genotypes can still trump matching the environment since the benefits from matching genotypes are positively frequency dependent and the penalty from

#### When matching the environment is much more advantageous than matching genotypes for only one partner (GXE >> GXG for only one partner)

When GXE interactions are strong for one partner (figure 1e-f) but weaker for the other (figure 1a-d), the only stable equilibrium is *migration-selection balance* (figure S2). This suggests that asymmetry in GXG vs. GXE interactions between mutualistic partners promotes genetic variation.

Because one set of genotypes derives a greater fitness advantage from matching genotypes but its partner derives a greater advantage from matching the environment, the *global genotypic match* or *mismatch*, where the same host/symbiont genotypes fix across all environments in parallel fashion for both partners, cannot be stable. The *reverse migration-selection balance* equilibrium, which relies on frequency dependence in the benefit from mutualism for both partners is not biologically relevant. This is because the benefits from mutualism exceed the benefit from matching environment for only one partner.

Despite our analysis of the internal equilibria assuming weak dispersal, simulations suggest that the *migration-selection balance* equilibrium remains the only locally stable equilibrium at high dispersal (figure S2).

#### When matching the environment is much more advantageous than matching genotypes (GXE >> GXG)

When GXE interactions are strong and GXG interactions are weak (figure 1e-f) in both partners, the *migration-selection balance* equilibrium is locally stable at high and low dispersal and the *global genotypic mismatch* equilibria are locally stable when dispersal is sufficiently large (figure 3). The stability of these equilibria suggest that strong GXE interactions can promote local genetic variation in mutualistic partners when dispersal is rare or erode it when dispersal is common.

**Figure 3:**
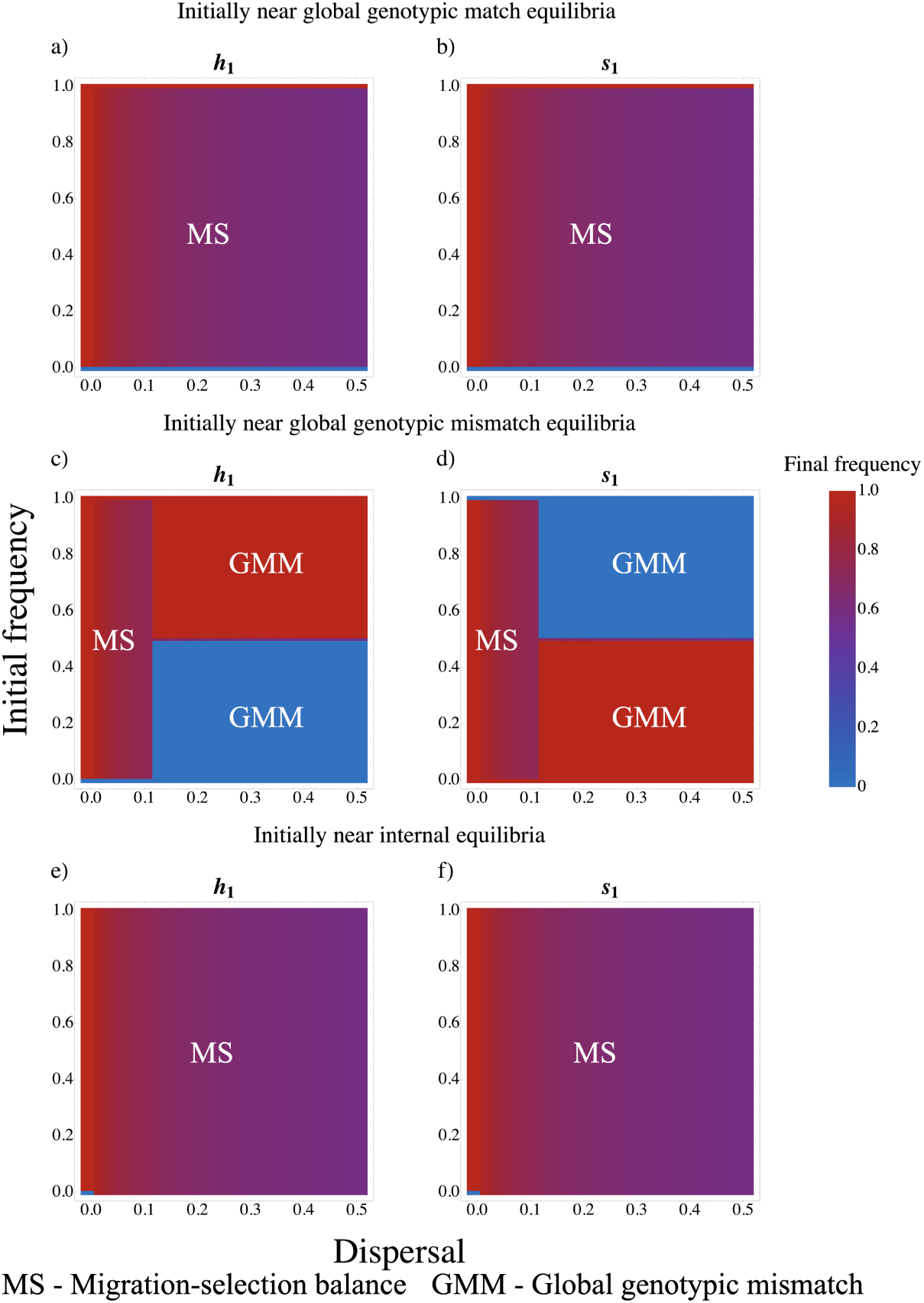
Results of simulations where matching the environment has a much greater effect on fitness than matching genotype (GXE *>>* GXG). Here Δ*H_ms_* = Δ*S_mh_* = 0.1 and Δ*H_gh_* = Δ*S_gs_* = −0.7. Each panel gives the frequency of genotype 1 in environment 1, *h*_1_ and *s*_1_ for hosts and symbionts, respectively, at *t* = 10, 000. The *x* axis is dispersal rate, *d*, and the *y* axis is the initial frequency of host genotype 1 in environment 1, *h*_1_. The other initial frequencies differ by row. In the first row, the initial frequencies of genotype 1 increase uniformly across both environments and species, *h*_1_ = *h*_2_ = *s*_1_ = *s*_2_. In the second row, as the initial frequencies of genotype 1 in the host increase the initial frequencies in the symbiont decrease, *h*_1_ = *h*_2_ = 1 − *s*_1_ = 1 − *s*_2_. In the third row, as the initial frequencies of genotype 1 in environment 1 increase the initial frequencies in environment 2 decrease, *h*_1_ = 1 − *h*_2_ = *s*_1_ = 1 − *s*_2_.

The *migration-selection balance* equilibrium is always locally stable under the small dispersal assumption. For low levels of dispersal, local selection favors the host and symbiont pairs that match both their environment and their partner over those that do not. *Migration-selection balance* also appears locally stable at high rates of dispersal (figure 3e-f), as matching both a partner and the environment remains advantageous.

As dispersal increases, the *global genotypic mismatch* equilibria also become locally stable (figure 3c-d). The mismatch equilibria become stable because, when dispersal is high enough to buffer against a local fitness disadvantage in the mismatched environment (figure S3a), a global fitness advantage is conferred as a host’s preferred symbiont becomes rare in both environments (figure S3b). This advantage arises because it allows the host to pair with an environmentally-matched symbiont in its mismatched environment while preventing the other host genotype from doing the same.

#### When hosts and symbionts have different dispersal rates

We next allowed hosts and symbionts to disperse at distinct rates, *d_h_* and *d_s_*, respectively. This modification recapitulated the general behaviors of the base model but exhibited a few key novel characteristics that suggest that asymmetric dispersal rates increase genetic variation.

When hosts and symbionts have different dispersal rates, the same four equilibria can be biologically valid, and their stability depends on GXG and GXE interactions in a similar fashion. However, the boundary equilibria, *global genotypic match* and *global genotypic mismatch*, differ in their stability criteria. Now the boundary equilibria increase in stability only if both *d_h_* and *d_s_* increase simultaneously (section S2). This more stringent criterion suggests that asymmetry in dispersal rates reduces the likelihood that genetic variation in mutualistic partners will be eroded. The model with asymmetric dispersal has another notable behavior. When the dispersal rate of one partner is low and GXE interactions are much stronger than GXG interactions, there is a novel locally stable equilibrium. At this equilibrium, the partner with a high dispersal rate fixes one genotype, while the other partner has a polymorphism (figure S4 top row, where the host fixes). This equilibrium occurs when the *global genotypic mismatch* would otherwise occur (figure 3 top row); the global genotypic mismatch is not realized because the slowly dispersing partner cannot wholly translate the benefits of matching the environment into fixation.

### Case studies

To demonstrate how to make predictions with our model and to examine the potential for GXE interactions to maintain genetic variation in mutualisms in nature, we next parametrize our model using data from three case studies. The results are summarized in Table 2.

**Table 2:** Summary of findings from case studies.

| System | Species | Strength of<br>GXE vs GXG<br>in hosts | Strength of GXE<br>vs GXG in<br>symbionts | Potentially<br>stable equilibria |
| --- | --- | --- | --- | --- |
| Plant-microbe<br>Ricks et al.<br>(2023) | *Bromus tectorum* and<br>microbiome | GXE >> GXG | Unknown | *Migration-*<br>*selection balance*<br>*and Global*<br>*genotypic*<br>*mismatch* |
| Plant-AM<br>fungi Johnson<br>et al. (2010) | *Andropogon gerardii*<br>and arbuscular<br>mycorrhizal fungi | GXE ≈ GXG | GXE > GXG | *Global genotypic*<br>*match and*<br>*Migration-*<br>*selection balance* |
| Fly-microbe<br>Henry et al.<br>(2020) | *Drosophila*<br>*melanogaster* and<br>*Acetobacter* spp. | GXE < GXG | Unknown | *All except Global*<br>*genotypic*<br>*mismatch* |

The first case study looks at environments likely connected by dispersal, while the second case study looks at environments that are far apart and the third is an experiment without any dispersal. In the latter two studies we are therefore examining the relative strength of GXE interactions and either assuming rare long-distance dispersal or hypothesizing about what would happen if dispersal was initiated.

For the two observational studies, we assume that the researchers sampled host and symbiont ecotypes from their locally adapted ‘home’ environments, and not ecotypes introduced from other environments via dispersal (in terms of our model, all hosts and symbionts sampled from environment *i* have genotype *i*). This essentially assumes the systems are near the *migration-selection balance* equilibrium with rare dispersal. If they were instead near the *global genotypic match* or *global genotypic mismatch* equilibria then we would not see any difference between hosts and symbionts from the two environments. And if they were instead near the *reverse migration-selection* equilibria then we would see host-symbiont pairs do (slightly) better in the environment they were not sampled from (*H_m_ < H_g_* and *S_m_ < S_g_*). Given the results below, we are therefore comfortable with our assumption that we are sampling locally-adapted genotypes.

#### Strong GXE interactions in a plant-microbe interaction

Ricks et al. (2023) conducted a reciprocal transplant experiment that measured the fitness of *Bromus tectorum* plant hosts collected from several highly saline and nearby non-saline environments with microbiomes that were sourced from the same highly saline or non-saline environments. Here we leverage their fully factorial experiment to parameterize our model’s payoffs and payoff differences (figure S5).

Natural systems will not display the symmetry in fitness payoffs we assumed. However, the plant hosts exhibit strong GXE interactions between environments (Δ*H_ms_*+ Δ*H_gh_ <* 0, GXE *>>* GXG) with both symbionts. Therefore, according to our model, the only equilibria that can be stable are the *global genotypic mismatch* and *migration-selection balance* equilibria, irrespective of the strength of GXG interactions between *B. tectorum* plants and their microbiomes, since strong GXE interactions trump strong GXG interactions in the other partner (see table 1). In other words, there is strong scope for GXE interactions to maintain variation within environments via *migration-selection balance* in this plant-microbe interaction.

Note that this plant-microbiome interaction is not a mutualism as plants did not benefit from microbial inoculation and had higher fitness with allopatric than with sympatric microbes (Ricks et al., 2023). However, this does not violate the assumption we made when assessing stability, that both Δ*H_ms_* and Δ*S_gh_* are positive. This case study therefore demonstrates that our model is general enough to also predict evolutionary dynamics in interactions that exhibit selection for antagonism.

#### Moderate GXE interactions in a plant-mycorrhizae mutualism

We next parameterize our model using Johnson et al. (2010)’s reciprocal transplant of *Andropogon gerardii* ecotypes and their local arbuscular mycorrhizal (AM) fungi among nutrient-limited grassland habitats. In this case the authors measured fitness for both plant hosts and AM fungi. We use the data from their transplant between Cedar Creek and Konza prairie to parameterize all the payoffs of our model.

For plant hosts (figure S6), there are no significant payoff differentials due to GXG or GXE interactions across both partners and environments. For AM fungi (figure 4), the strength of GXE interactions is asymmetric depending on which host genotype the symbionts are partnered with. When partnered with the Konza host ecotype, GXE interactions are moderate (Δ*S_gs_ <* 0 but Δ*S_mh_* + Δ*S_gs_ >* 0, figure 4b), but when partnered with the Cedar Creek host ecotype, GXE interactions are strong (Δ*S_mh_* + Δ*S_gs_ <* 0, figure 4d).

**Figure 4:**
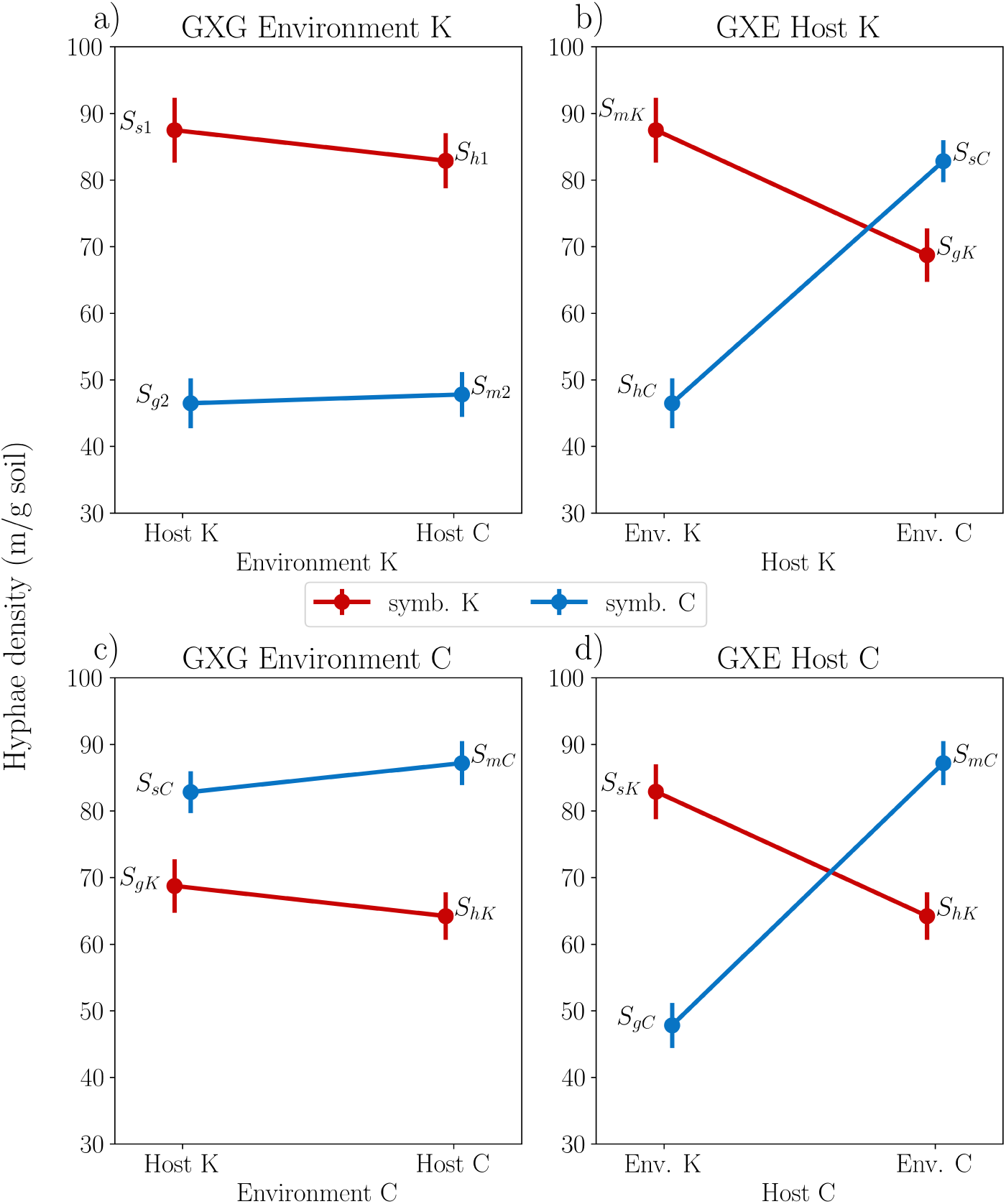
Empirical strengths of GXE and GXG interactions for arbuscular mycorrhizal (AM) fungi in plant-AM fungi symbiosis (Johnson et al., 2010). Because in the real world, environments and partners are not perfectly symmetrical, there are plots for the strength of the GXG interactions in each environment (panels a and c) and the GXE interactions across each symbiont type (panels b and d). On the y axis we use Johnson et al. (2010)’s proxy for AM fungal fitness: hyphae density. *K* denotes a Konza prairie environment or genotype and *C* denotes a Cedar Creek environment or genotype. Points are labeled with their corresponding payoff by symbiont ecotype (e.g., *S_gK_* is symbiont *K*’s payoff from matching host *K* and *S_sK_* is symbiont *K*’s payoff from matching environment *K*). Error bars are standard errors of the means.

Because the data violate our assumption that the strength of GXE interactions is symmetric between partners and environments, we simulate the behavior of an asymmetric version of the model (section S3) with the parameter values from this study. This new version of the model behaves similarly to the symmetric model when symbionts receive the mean payoff from each host type (figures S7-S10).

GXE interactions are moderate on average (Δ*S_gs_ <* 0 but Δ*S_mh_*+ Δ*S_gs_ >* 0, GXE > GXG), meaning that our prediction is that *migration-selection balance* will be locally stable for any level of dispersal (table 2), which is what we see in asymmetric simulations (figure S7). We also predict that the *global genotypic match* equilibria are locally stable with frequent dispersal. The only difference in the asymmetric simulations is that only one *global genotypic match* equilibrium, where the Konza AM fungi ecotype fixes globally, appears locally stable when dispersal is high (figure S9). This is because, in the model, the Konza ecotype experiences a lower fitness penalty for mismatching its environment (Δ*S_gs_ <* 0 but Δ*S_mh_* + Δ*S_gs_ >* 0) allowing it to fix globally with its preferred host. However, as dispersal is unlikely to be frequent between Konza and Cedar Creek, which are separated by approximately 750km, the most likely stable equilibrium is *migration-selection balance*.

#### Strong GXG in the Drosophila melanogaster microbiome

Finally, Henry et al. (2020) conducted a microbiome reciprocal transplant experiment that we use to parameterize our fitness payoffs. They experimentally evolved one population of *Drosophila melanogaster* together with its microbiome (composed of *Acetobacter* spp.) on a stressful, high sugar (HS) diet. They then reciprocally transplanted microbiomes between control (C) flies evolved on a normal diet and flies evolved on the HS diet and measured fly reproductive output under C and HS conditions. The resulting fitness payoffs (figure S11) indicate strong GXG interactions for host fitness across both environments (Δ*H_ms_* & Δ*H_gh_ >* 0, GXE < GXG). Without knowing the strength of GXG and GXE interactions in the microbiome, the most we can say is that this rules out the *global genotypic mismatch* equilibria. More data is needed to predict how likely it is that GXE interactions could maintain polymorphism in partner quality via *(reverse) migration-selection balance* in this system.

## Discussion

Our analysis finds four scenarios that reflect the impact of increasing the strength of GXE interactions on mutualistic coevolution and the resulting genetic variation (table 1). When GXE interactions are effectively absent (GXE < GXG, figure 1a-b), there is one non-polymorphic and at least one polymorphic stable equilibrium (figure 2). Variation in mutualisms can be maintained in the absence of GXE interactions by strong positive frequency-dependence and spatial structure. Increasing the strength of GXE interactions so that they are just larger than GXG interactions (GXE > GXG for one or both partners, figure 1c-d), the non-polymorphic equilibrium loses stability at low dispersal (figure S1), promoting variation. Making GXE interactions much stronger than GXG interactions for one species (figure 1e-f) takes this one step further, guaranteeing locally stable polymorphism via migration-selection balance at any dispersal rate (figure S2). Interestingly, when both species have GXE *>>* GXG, the possibility of no polymorphism returns, but only for large dispersal rates (figure 3), where a genotype can gain a global fitness advantage when its preferred partner is absent (figure S3).

### Implications for empirical studies

Our theoretical framework, when parameterized with existing empirical data from local adaptation studies, predicts the amount of genetic variation maintained by GXE interactions and the mechanism by which they do so. Many mutualism researchers conduct reciprocal transplant studies in order to understand the extent to which mutualistic interactions mediate adaptation to different environmental conditions (Petipas et al., 2021). When these studies test fitness in a fully factorial fashion across mutualistic partners and environments, they can be used to parameterize the fitness payoffs of our model, as we have shown for three case studies. These types of reciprocal adaptation studies, especially when paired with estimates of dispersal between environments, may provide important insights into the process of coevolution between mutualists, and the amount of variation that is expected to be maintained. Two of the three examples here suggest moderate to strong GXE interactions that can maintain substantial variation.

In addition to parameterizing our payoffs, the fitnesses of mutualistic partners in a fully factorial reciprocal transplant study could indicate which equilibrium the system is near. Populations near the *global genotypic match* or *global genotypic mismatch* equilibria fix a single host genotype and a single symbiont genotype in both environments, so a reciprocal transplant would find null results (i.e., no GXG or GXE interactions). In contrast, near the *reverse migration-selection balance* equilibria, there are distinct frequent host and symbiont ecotypes in each environment and a reciprocal transplant would find that hosts and symbionts perform better in their ‘away’ environment than in their ‘home’ environment. However, since the *reverse migration-selection balance* equilibria are stable and valid only when GXE interactions are relatively weak or absent, it is possible that host and symbiont fitnesses would not vary much or at all between environments (i.e., no GXE interactions). Finally, if the populations are near *migration-selection balance*, hosts and symbionts will have distinct ecotypes that have the highest fitness in their sympatric environment and with their sympatric partner (both GXE and GXG interactions). As *migration-selection balance* is the most likely stable equilibrium when dispersal is rare or there are there are asymmetries in the relative strength of GXE across partners (table 1), these two conditions increase the chances that an experiment detects GXG and GXE interactions.

Our model’s findings also provide an explanation for some non-intuitive results from studies of mutualisms in the wild. There is empirical evidence for local maladaptation between mutualistic partners in some systems (Gómez et al., 2009; Johnson et al., 2010; Rúa et al., 2016). For example, in the symbiosis between corals and their algal photosymbionts, many corals harbor *S. trenchii* symbiodinium that are well adapted to their environment, but maladapted to their hosts (Pettay et al., 2015). This result could be explained by local selection for parasitism; however, our model provides another explanation: strong GXE and high dispersal can lead to a stable *global genotypic mismatch*. Certain combinations of mutualists can also appear maladapted to their environment (Rúa et al., 2016). This could be due to coevolution between mutualists slowing the approach to a stable environmentally matched equilibrium (Thompson et al., 2002); however, our model suggests that this scenario could also be a stable *reverse migration-selection balance* equilibrium if GXE interactions are relatively weak and dispersal is low (e.g., as in the *Drosophila melanagoster* microbiome experiment, Henry et al., 2020). Finally, our model suggests that asymmetric rates of dispersal between partners increases the scope for local polymorphism in mutualisms. This scenario is likely in many partnerships in which both partners have distinct ecological characteristics. For example, even in a partnership in which one species has a high rate of dispersal, a rate-limiting mutualistic partner may still promote variation. This could be the case in many plant-microbe mutualisms, where microbial dispersal rates are largely unknown (Baltrus, 2020). Taken together, these findings may help to explain observations from studies of local adaptation and local genetic variation in mutualism.

### Implications for mutualism theory

Our work builds on classic results from coevolutionary theory, a field that has long understood that in the absence of spatial structure, mutualisms mediated by reciprocally beneficial matching alleles tend to lose genetic variation (Nuismer, 2017, chapter 2). To our knowledge, there is only one published model of mutualism where the mutualistic interaction is reciprocally beneficial but environmentally contingent (Nuismer, 2017, chapter 8). This model assumes that in a mismatch between genotypes, mutualistic partners receive no benefit, and thus only considers the GXE < GXG scenario. To illustrate, setting *H_s_* = 0 and *H_h_* = 0 in figure 1, results in GXG > GXE interactions. When GXE < GXG, our model predicts that three equilibria can be valid and locally stable (table 1). However, Nuismer (2017, chapter 8) finds that only the *global genotypic match* equilibria are stable. Why does the Nuismer (2017) model draw such a different conclusion than ours? The chief reason for the difference is that the Nuismer (2017) model assumes selection is weak relative to dispersal. If this model is instead analyzed with weak dispersal and strong selection, both internal equilibria are stable (see nuismer.nb in our online supplement). Our analysis thus adds to our theoretical understanding of coevolution between mutualists by suggesting that even when mutualists experience mutually beneficial GXG interactions, it is still possible for GXE interactions to maintain genetic variation under certain, previously unexplored, conditions.

The results of our model provide novel insights into coevolution between mutualists while also re-affirming classic general theoretical expectations. Our model is consistent with the geographic mosaic theory of coevolution (Thompson, 2005) because we assume that the benefit from mutualism depends on both the match between partner genotypes and their environment; thus, our model involves a genotype-by-genotype-by-environment (GXGXE) interaction for fitness payoffs (figure S.12a-c). However, because we assume that fitness payoffs are symmetric between environments, there is no GXGXE interaction for relative fitness in our model (figure S.12d-f) and therefore no effect of GXGXE on the dynamics. Our simulations in our plant-AM fungi case study suggest that a model that relaxes this symmetry assumption exhibits similar characteristics to our symmetric model, but analytical work along these lines is still needed.

Spatial structure can maintain variation in mutualistic interactions with mutually beneficial GXG interactions (Carlson et al., 2022); however, our model suggests that strong benefits from matching genotypes may at times trump weak to moderate GXE interactions for fitness (first row 1). And even in the absence of strong frequency-dependent selection due to GXG interactions, we found that variation may be eroded due to strong GXE interactions, leading mutualistic partners to be maladapted to one another when dispersal is high (*global genotypic mismatch*). That coevolution generates maladaptation is not a new insight (Thompson et al., 2002); however, our model presents a novel mechanism by which this maladaptation might occur – mismatched partners are stable because they help each other in their respective environments.

Many mutualisms are subject to a ‘new’ paradox. Mutualisms are thought to be maintained via costly stabilizing traits such as partner choice, sanctioning, and screening that improve the relative fitness of cooperative mutualistic partners. However, these traits erode their own selective incentive: variation in mutualistic partner quality (Heath and Stinchcombe, 2014). Our model explores the extent to which one proposed source of variation in mutualisms, GXE interactions, maintains variation in two interacting partners. Our results suggest that GXE interactions could stably maintain variation in mutualism upon which stabilizing traits might act. We anticipate that incorporating partner choice or vertical transmission into our model would likely reduce the scope for spatial structure to maintain variation within environments when GXE interactions are relatively weak (by destabilizing *reverse migration selection balance*) but increase local genetic variation when GXE interactions are relatively strong (by destabilizing the *global genotypic mismatch* equilibria). Synchronous dispersal of pairs of hosts and symbionts would likely have the same effect. These intuitions suggest that a full analysis examining how the evolution of mutualism-stabilizing traits is influenced by GXE interactions is an intriguing avenue for future research.

## Conclusion

Understanding the mechanisms that maintain variation in partner quality is a key question for mutualism researchers given that mutualists likely experience strong reciprocal selection when GXG interactions are mutually beneficial (Heath and Stinchcombe, 2014). Here, we have shown that GXE interactions have the potential to maintain variation in partner quality in mutualisms even if mutualistic partners do not experience selection to become antagonistic. GXE interactions have long been known to cause variation in traits (Muir et al., 1992); however, our work suggests that, in mutualisms, the magnitude of GXE interactions relative to GXG interactions can cause GXE interactions to maintain – or erode – genetic variation in complex and at times counter-intuitive ways.

## Acknowledgements

The authors thank Nathaniel Carlson, Julia Boyle, and Osmond lab for helpful reviews of the manuscript and the Frederickson and Stinchcombe labs for helpful feedback and discussions. We also thank Kevin Ricks, Nancy Collins Johnson, Gail Wilson, and Lucas Henry for contributing data for case studies.

## Author Contributions

All authors contributed to the conception of the study and the final manuscript. C.C. and M.O. carried out mathematical analysis of the model. C.C. wrote the first draft of the manuscript, analyzed numerical simulations, and made the figures.

## Data availability

The code for the stability analysis and numerical simulations were written in Mathematica and can be found at https://github.com/chrisiancarlson/GXE-mutualism.

## Supplementary material

### S.1 Fitnesses

Let *w_ji_*(*s_i_*) denote the fitness of host genotype *j* in environment *i* as a function of symbiont 1’s frequency in environment *i*. Our expressions for host fitnesses are:

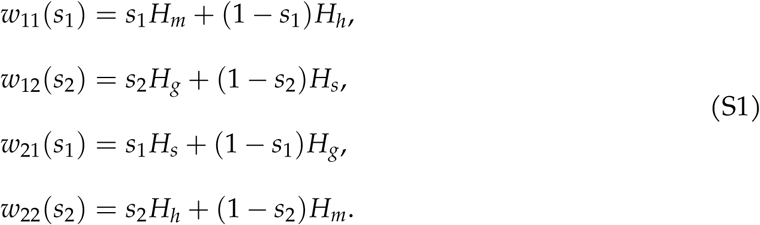

Similarly, let *v_ji_*(*s_i_*) denote the fitness of symbiont genotype *j* in environment *i* as a function of the frequency of host genotype 1 in environment *i*. Our expressions for symbiont fitnesses are:

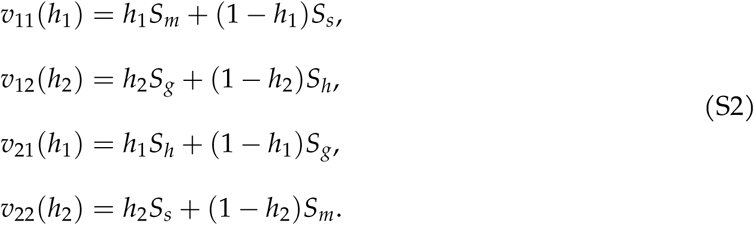

### S.2 Equilibria and their stability

Here we present a more general analysis, allowing hosts and symbionts to have distinct dispersal rates, *d_h_* and *d_s_*, respectively.

The local stability of an equilibrium of equation 3, (*h*_1_, *h*_2_, *s*_1_, *s*_2_), is determined by the eigenvalues of the Jacobian

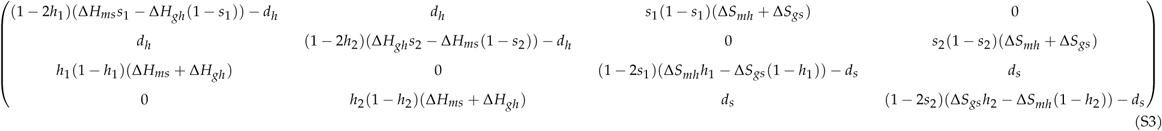

evaulated at that equilibrium.

From equation 3, we first numerically found all boundary equilibria, where the frequency of host and symbiont genotypes in either environment is 0 or 1. The Jacobian at these equilibria is block diagonal, providing us with relatively simple Routh-Hurwitz local stability criteria (Otto and Day, 2011, box 8.2).

We next found and approximated the internal equilibria using a first order Taylor series expansion of equation 3 assuming that dispersal is small. We then evaluated the local stability of the approximate equilibria by approximating the characteristic polynomial of the Jacobian evaluated at the equilibrium of interest and solved this expression for its eigenvalues (Otto and Day, 2011, section 8.4).

When assessing stability, we assume that Δ*H_ms_* and Δ*S_mh_* will always be positive as a full match should provide the largest benefit. The other two payoff differences, Δ*H_gh_* and Δ*S_gs_*, can be positive or negative, depending on the value of the payoff derived from a genotypic match (GXG interaction) vs. the payoff derived from a genotype matching with its environment (GXE interaction).

#### S.2.1 Global genotypic match

There are two *global genotypic match* equilibria, one where host 1 and symbiont 1 fix in both environments, (1, 1, 1, 1), and another where host 2 and symbiont 2 fix in both environments, (0, 0, 0, 0). The Routh-Hurwitz criteria for the stability of both of these *global genotypic match* equilibria are:

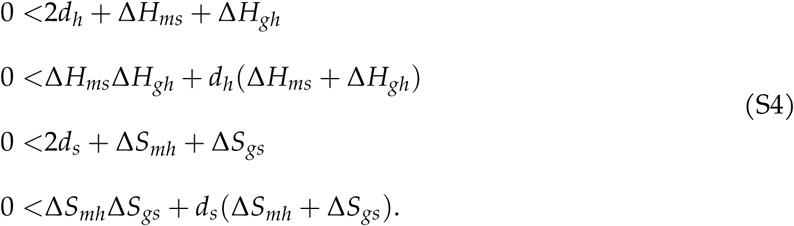

The local stability of this equilibrium depends on two quantities in each partner, the sum (e.g., Δ*H_ms_*+ Δ*H_gh_*for the host) and product (e.g., Δ*H_ms_*Δ*H_gh_* for the host) of payoff differences. These correspond to the arithmetic and geometric mean fitness advantage of the genotype near fixation. When dispersal is rare, the geometric mean is most important as stability requires that the dominant genotype will increase in frequency in both environments, e.g., Δ*H_ms_*Δ*H_gh_ >* 0 for the host. Under our assumption, Δ*H_ms_ >* 0, this requires Δ*H_gh_ >* 0, which we can interpret at GXE>GXG. As dispersal increases, individuals experience both environments with more frequency, increasing the importance of the arithmetic mean. Stability therefore increases with dispersal when the arithmetic mean is positive, Δ*H_ms_*Δ*H_gh_ >* 0. At very high rates of dispersal we can consider the two environments as one and only the arithmetic mean matters.

#### S.2.2 Global genotypic mismatch

The model also has two *global genotypic mismatch* equilibria, where a host genotype fixes with its mismatched symbiont genotype (i.e., host 1 and symbiont 2 fix in both environments, or host 2 and symbiont 1 fix in both environments). The stability criteria for these two equilibria are:

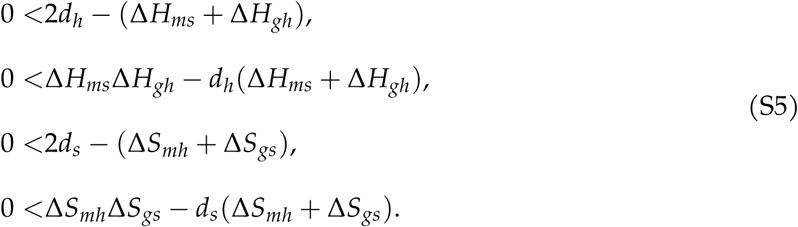

As with the *global genotypic match*, stability depends entirely on the sum and product of payoff differences, which again correspond to the arithmetic (e.g., −(Δ*H_ms_*+ Δ*H_gh_*) for the host) and geometric (e.g., Δ*H_ms_*Δ*H_gh_* for the host) fitness advantage of the dominant genotypes. The same intuition follows, with the arithemtic mean advantage becoming more important as dispersal increases. However, when there is little dispersal this equilibrium cannot be stable because the dominant genotype cannot increase in frequency in its mismatched environment (e.g., −Δ*H_ms_ <* 0 for the host).

#### S.2.3 Migration-selection balance

One approximate internal equilibrium is *migration-selection balance*:

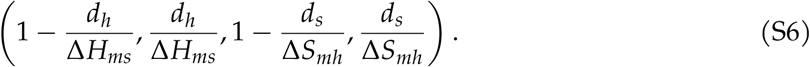

That is, host 1 and symbiont 1 are common in environment 1, and maintained at low frequency in environment 2 by dispersal.

The approximate stability criteria for this equilibrium are:

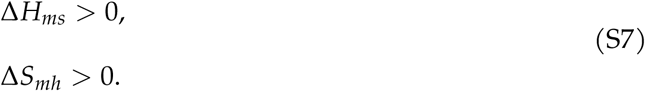

These conditions will always be true under our assumption that the matching payoff exceeds any mismatched payoff. As we approximate this equilibrium and its stability under rare dispersal, we use simulations to determine what happens as the *d_i_*approach the payoff differences.

#### S.2.4 Reverse migration-selection balance

Another approximate internal equilibrium is:

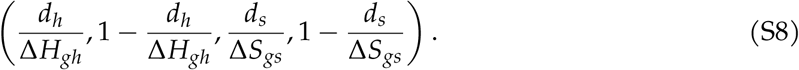

This is a mirror image of the *migration-selection balance* equilibrium, with host 1 and symbiont one now common in environment 2 and maintained in environment 1 by dispersal. We therefore call this equilibrium *reverse migration-selection balance*. It is only biologically valid when Δ*H_gh_ >* 0 and Δ*S_gs_ >* 0, i.e., when GXG>GXE.

The approximated stability criteria for this equilibrium are:

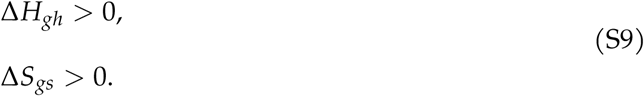

Therefore, this equilibrium is stable whenever it is biologically valid. Essentially, we need the positive feedbacks from GXG interactions to maintain a pair of matching genotypes in their mismatched environment. Again, it is not clear what happens to this equilibrium as dispersal gets large so we use simulations.

### S.3 Asymmetric model analysis

We used the fitness payoffs from Johnson et al. (2010) to parameterize our model. Because the payoffs of S6 did not conform to our symmetric model assumption, we simulated the behavior of an asymmetric version of our model, where payoffs can be different across environments. The expressions for the fitness of a focal host or symbiont in the asymmetric version of the model are:

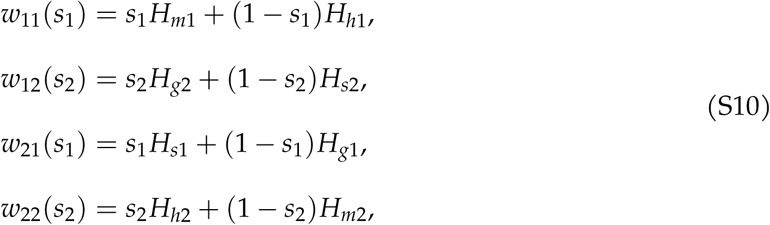

and

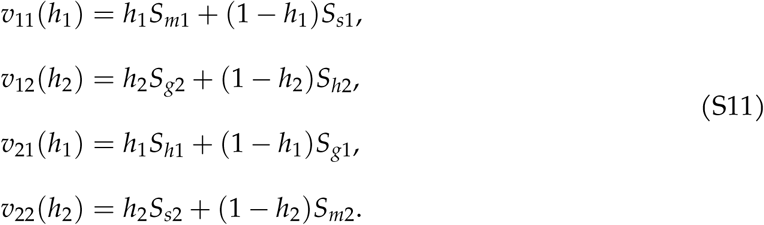

We simulated this asymmetric version of our model and include the results in our supplemental mathematica notebooks. We found that the asymmetric version of our model exhibited similar qualitative behaviors to the symmetric version when we averaged the values of the symbiont payoffs between partners, e.g., 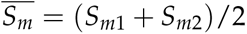.

### S.4 Supplementary figures

**Figure S1:**
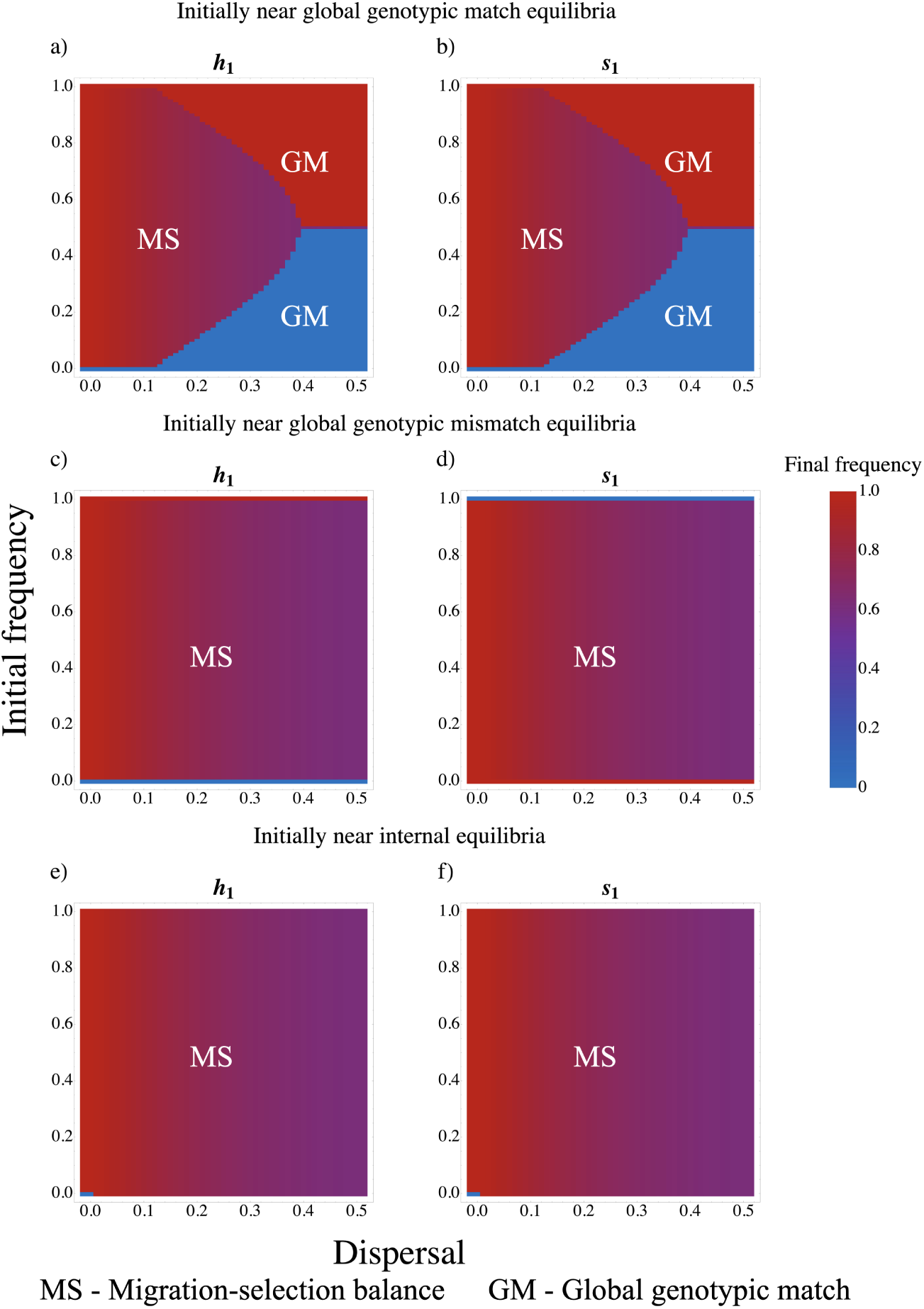
Results of simulations when GXE > GXG. Here Δ*H_ms_* = Δ*S_mh_* = 0.7 and Δ*H_gh_* = Δ*S_gs_* = −0.1. See figure 2 for more information.

**Figure S2:**
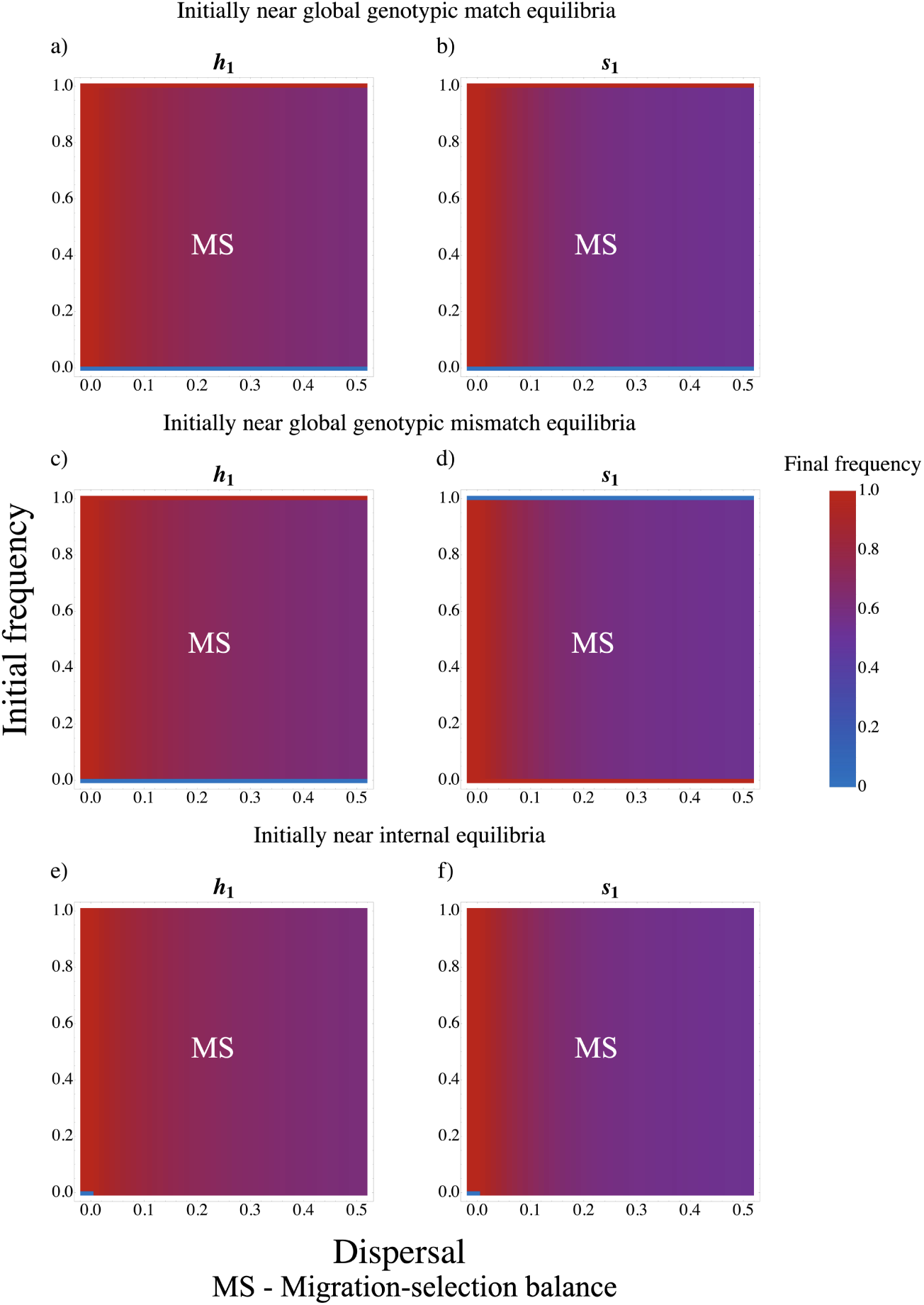
Results of simulations when GXE *>>* GXG for one partner and GXE < GXG for the other. Here Δ*H_ms_* = Δ*S_mh_* = 0.3, Δ*H_gh_* = −0.7, and Δ*S_gs_* = 0.1. See figure 2 for more information.

**Figure S3:**
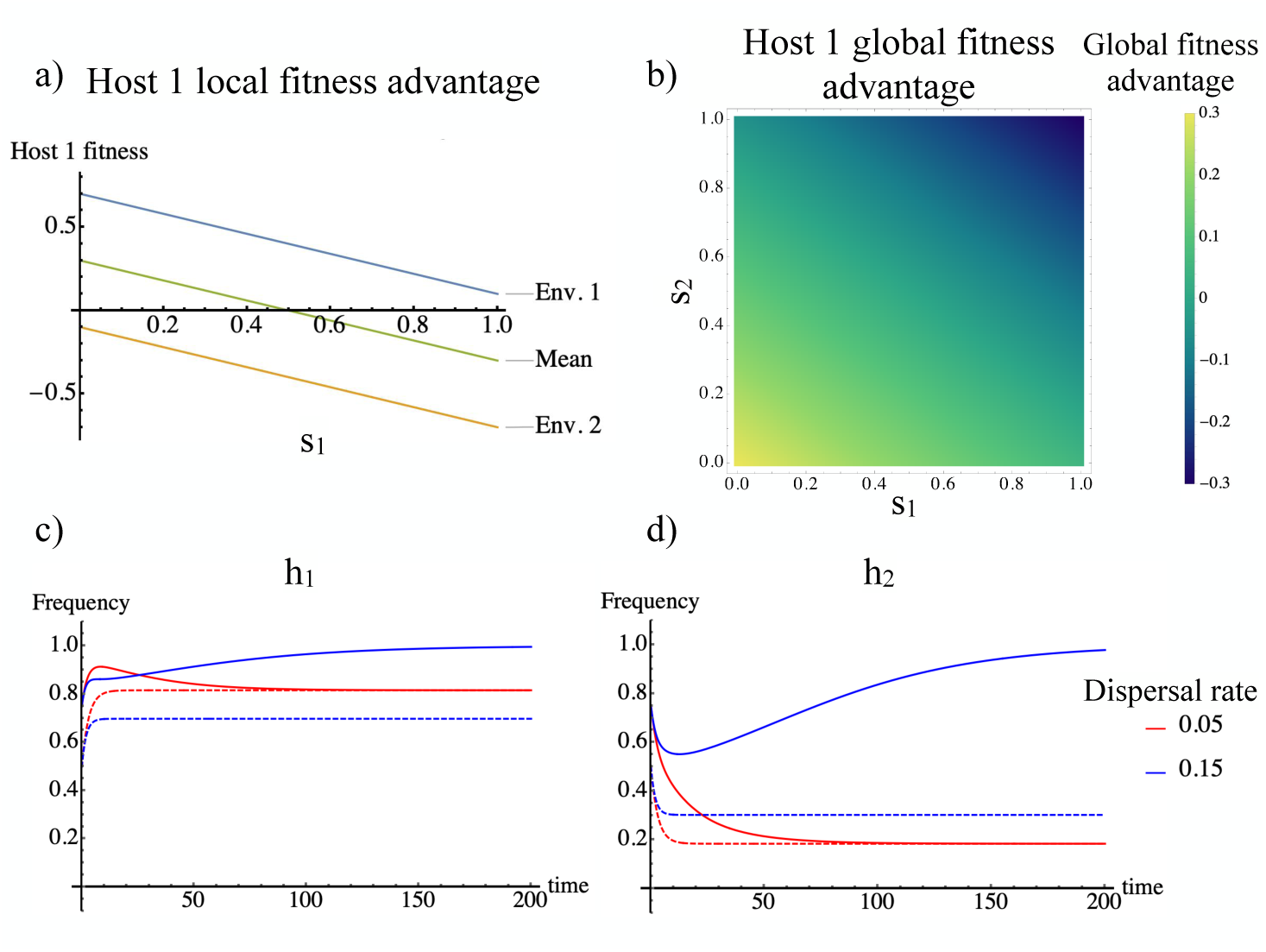
Fitness advantage for host 1 (*A* and *B*) in the case when matching environment has a greater effect on fitness than matching genotypes (GXE *>>* GXG) and the associated frequency dynamics (*C* and *D*). Here, Δ*H_ms_* = Δ*S_mh_* = 0.1 and Δ*H_gh_* = Δ*S_gs_* = −0.7. (*A*) The fitness advantage of host 1 in each environment (blue and orange), Δ*w_i_*(*s_i_*), and its average advantage across environments (green) as a function of the frequency of symbiont genotype 1, *s_i_* (the green line assumes the symbiont frequency is equal in the two environments, *s*_1_ = *s*_2_). (*B*) The average fitness advantage of genotype 1 across environments (color scale) as a function of the frequency of symbiont 1 in environment 1 (x-axis) and 2 (y-axis). (*C* and *D*) The dynamics of host 1 frequency when host initial frequency and dispersal vary and symbiont 1’s initial frequency is *s_i_* = 0.5 in both environments. In both plots, the initial frequency of host 1 is either *h_i_* = 0.5 (dashed) or *h_i_* = 0.75 (solid) in both environments. The red lines show host frequency when dispersal is low (*d* = 0.05) and the blue lines show host frequency when dispersal is high (*d* = 0.15).

**Figure S4:**
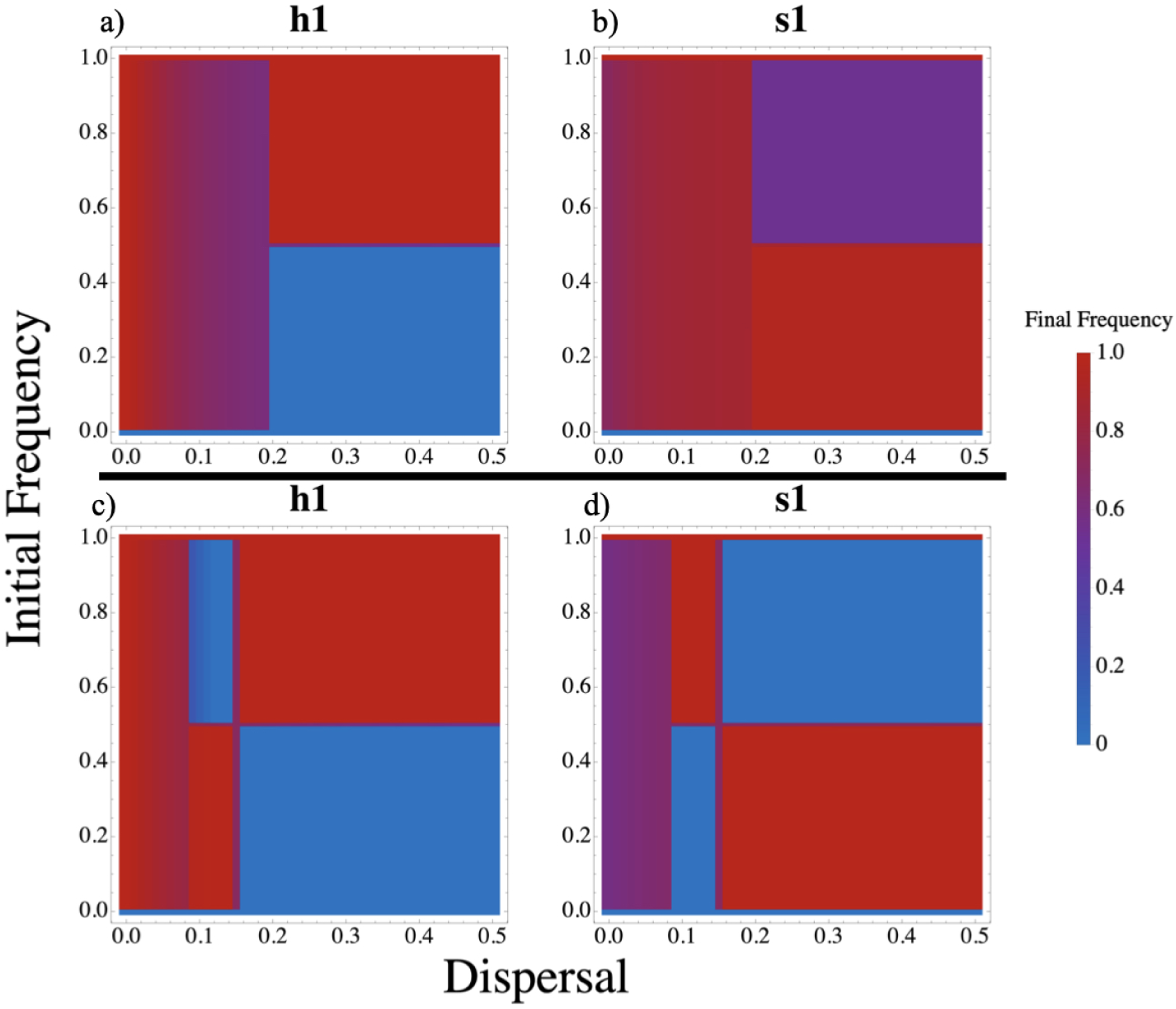
Results of simulations where matching environment has a much greater effect on fitness than matching partner genotype (GXE*>>*GXG). Here Δ*H_ms_* = Δ*S_mh_* = 0.1 and Δ*H_gh_* = Δ*S_gs_* = −0.7. The *x* axis is host dispersal rate, *d_h_*, and the *y* axis is the initial frequency of genotype 1 in hosts in both environments. In the top row, as the initial frequencies of genotype 1 in hosts increase the initial frequencies in symbionts decrease (*h*_1_ = *h*_2_ = 1 − *s*_1_ = 1 − *s*_2_) and the symbiont has a relatively low dispersal rate (*d_s_* = 0.05). In the bottom row, the initial frequencies of genotype 1 increase in synchrony (*h*_1_ = *h*_2_ = *s*_1_ = *s*_2_) and the symbiont has a relatively high dispersal rate (*d_s_* = 0.15). With symmetric dispersal (varying *d_s_* = *d_h_* on the x axis), the top panel would be composed of *migration selection balance* and *global genotypic mismatch* while the bottom panel would be only *migration selection balance* (figure 3 bottom row). Color shows the frequency of a given genotype in a given environment at *t* = 10, 000.

**Figure S5:**
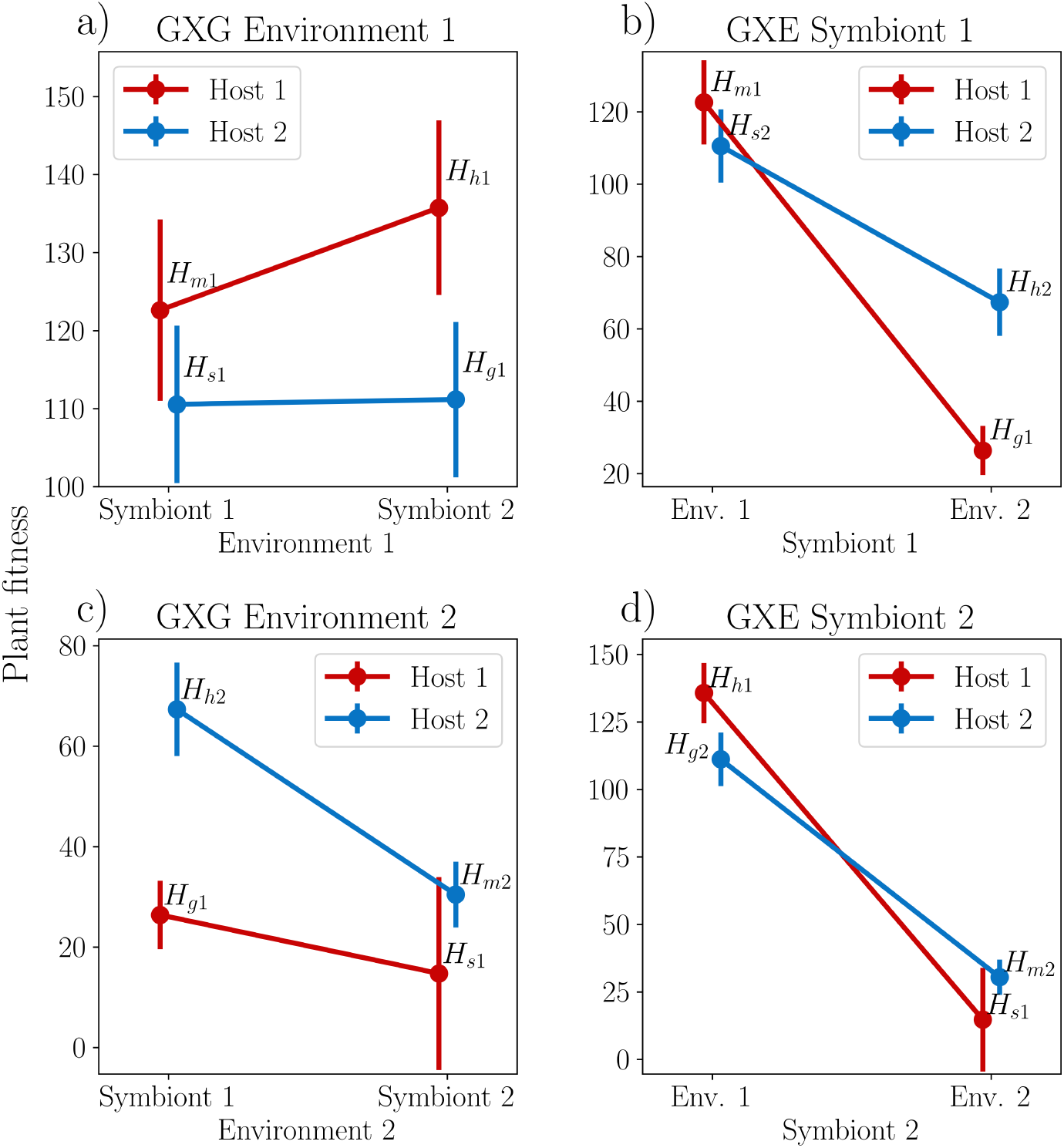
Empirical strengths of GXE and GXG in a plant microbe interaction (Ricks et al., 2023). Because in the real world environments and partners are not perfectly symmetrical, there are plots for the strength of GXG in each environment (a and c) and GXE across each symbiont type (b and d). 1 denotes a nonsaline environment and genotype and 2 denotes a saline environment and genotype. Points are labeled with their corresponding payoff (e.g. *H_h_*_1_ is host 1’s payoff from matching environment 1 and *H_h_*_2_ is host 2’s payoff from matching environment 2). Error bars are the standard error calculated by Ricks et al. (2023)

**Figure S6:**
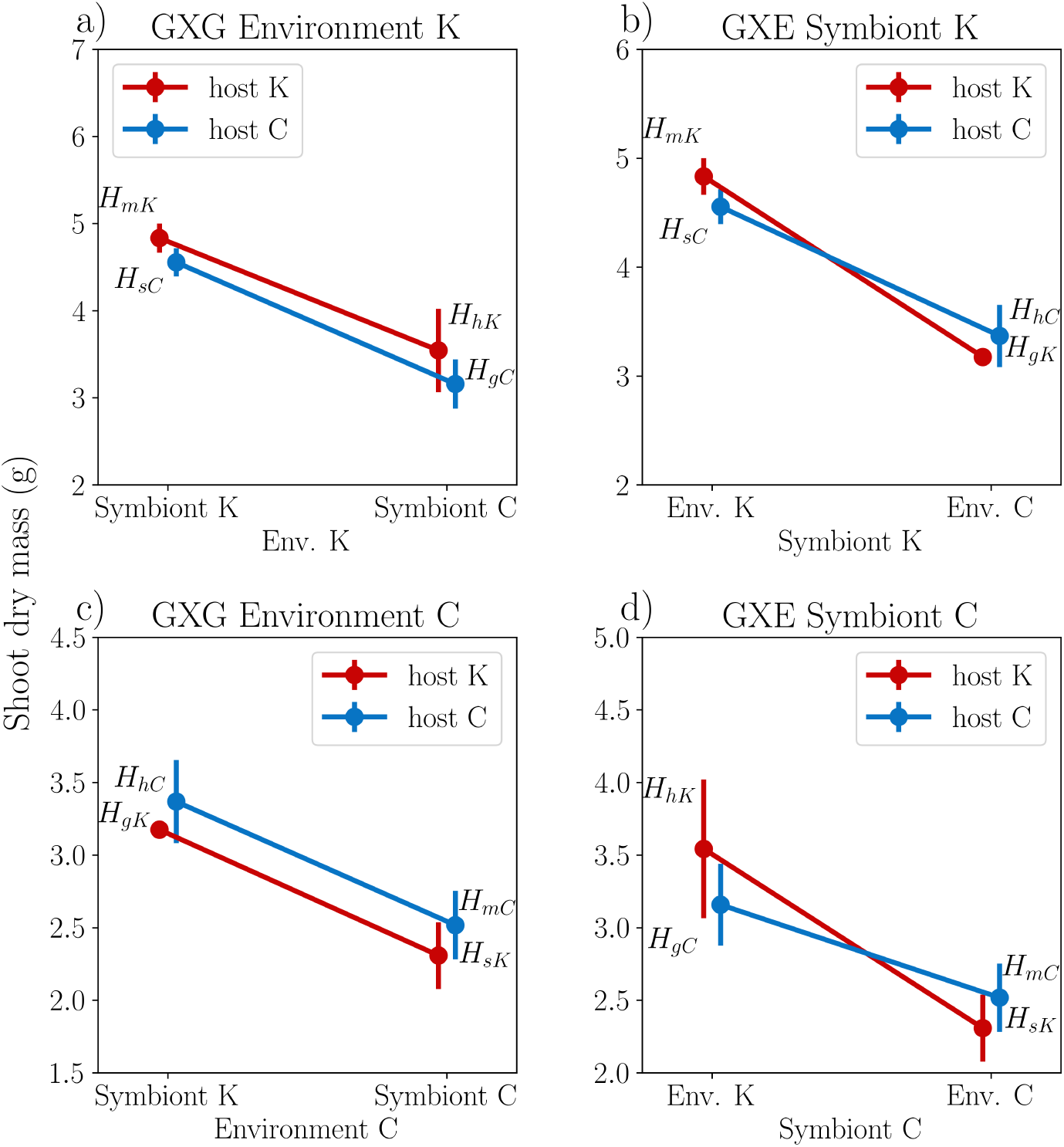
Empirical strengths of GXE and GXG for plants in plant-arbuscular mycorrhiza (AM) fungus mutualism (Johnson et al., 2010). *K* denotes a Konza Prarie environment or genotype and *C* denotes a Cedar Creek environment or genotype. Points are labeled with their corresponding payoff. Error bars are standard errors.

**Figure S7:**
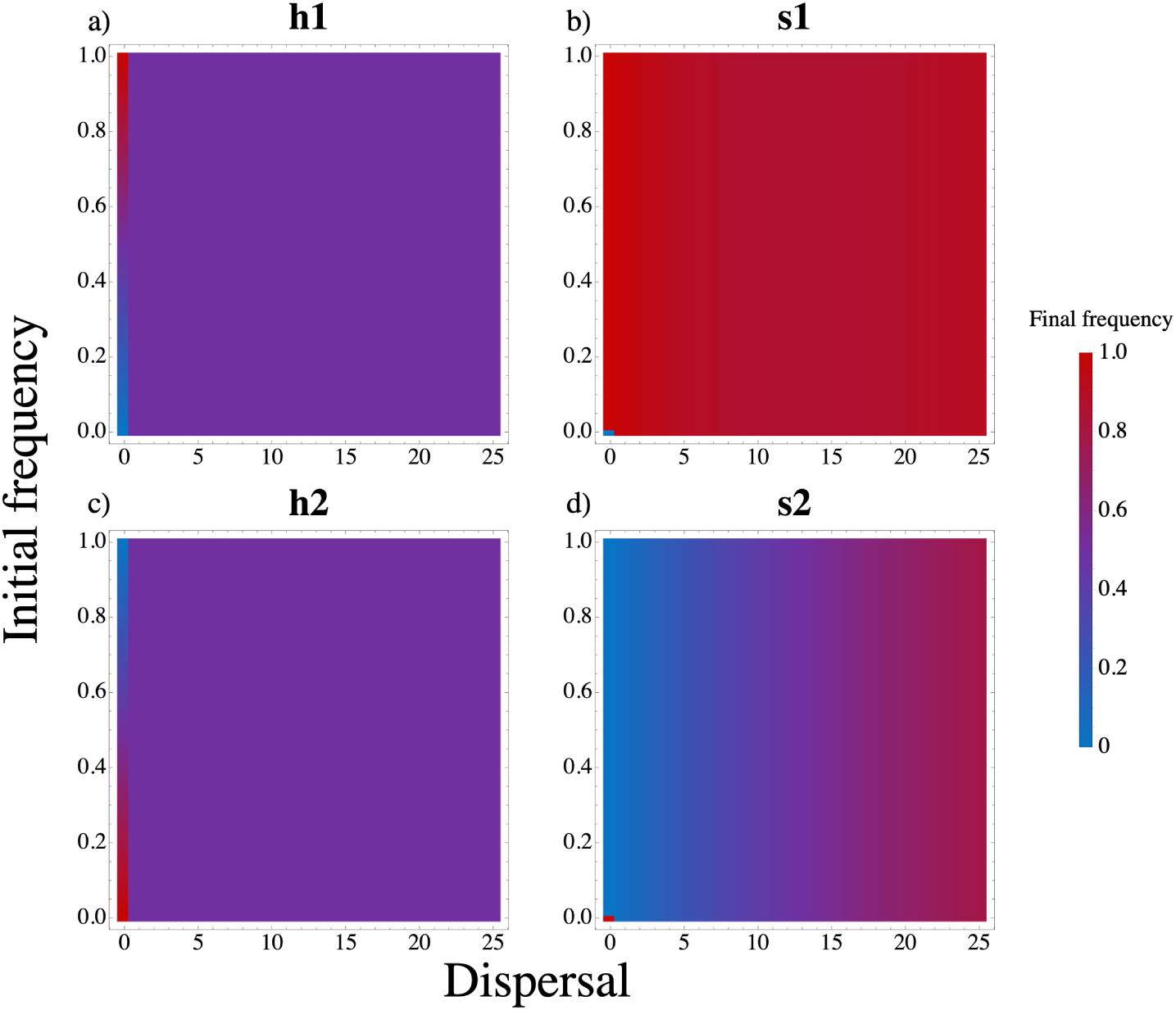
Results of simulations using asymmetric parameters from Johnson et al. (2010). Because there were no significant host payoff differentials we used the same average payoffs for all hosts. Each panel gives the final frequency of genotype 1 in a given species and environment at *t* = 10, 000. The *x* axis is dispersal rate, *d*, and the *y* axis is the initial frequency of genotype 1 in environment 1 in both hosts and symbionts. As the initial frequencies of genotype 1 in environment 1 increase the initial frequencies in environment 2 decrease, *h*_1_ = (1 − *h*_2_) = *s*_1_ = (1 − *s*_2_) (i.e., we are near the *migration-selection balance* equilibrium). 1 refers to Konza Prairie and 2 refers to Cedar Creek.

**Figure S8:**
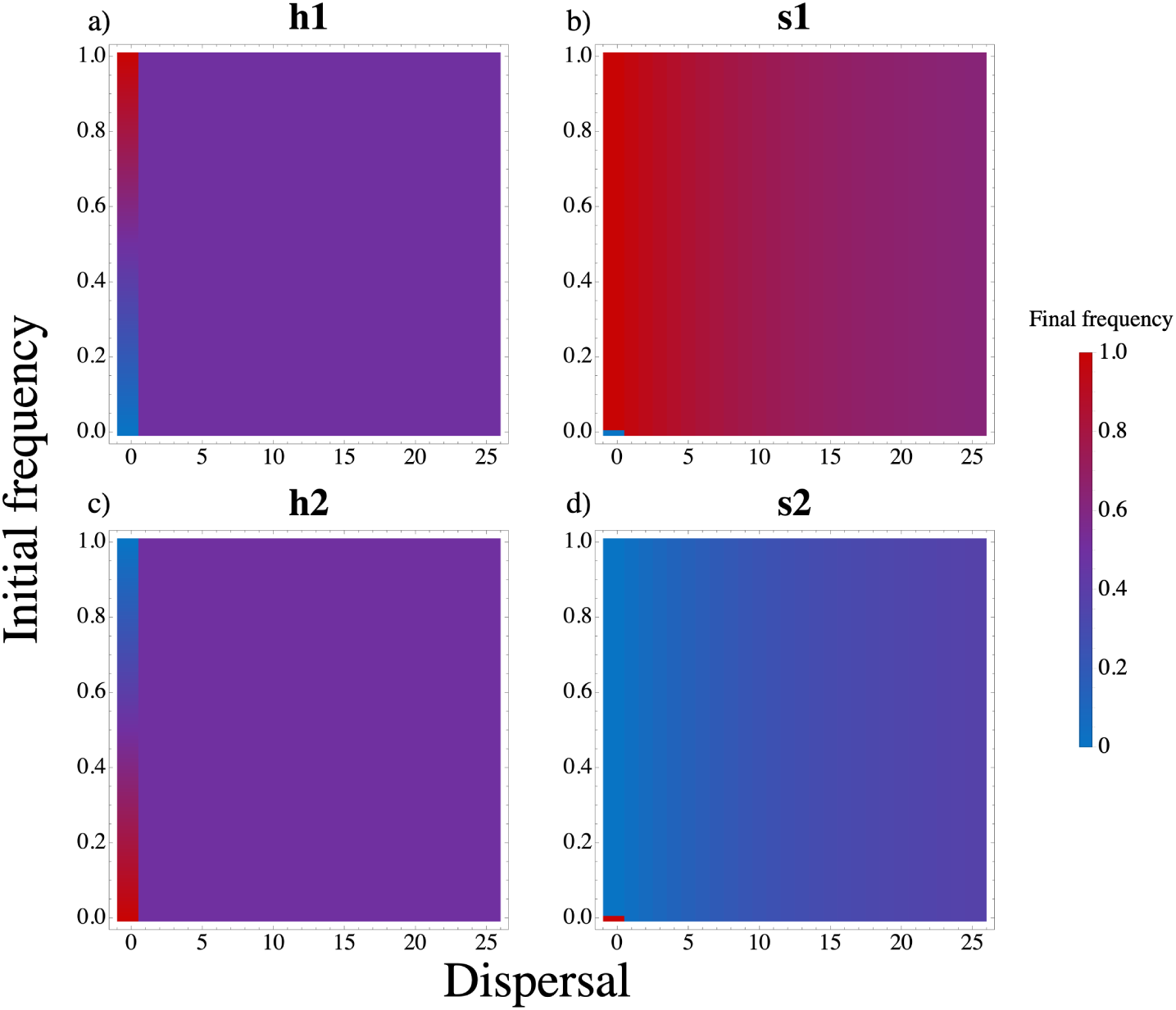
Results of simulations using parameters from Johnson et al. (2010), but averaged across each host and symbiont genotype. See figure S7 for more information.

**Figure S9:**
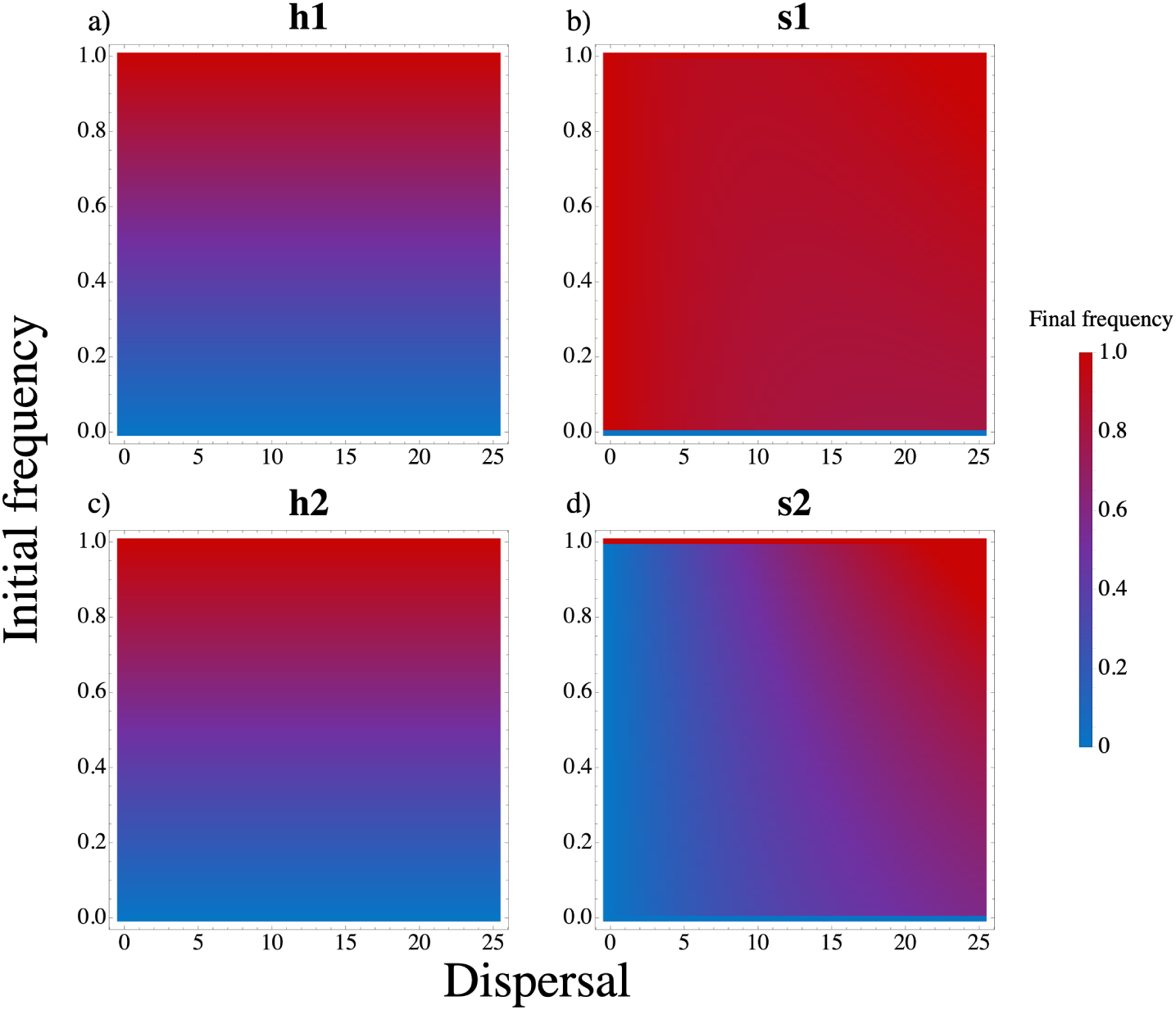
Results of simulations using, asymmetric parameters from Johnson et al. (2010). Because there were no significant host payoff differentials we used the same average payoffs for all hosts. Initial frequencies increases uniformly acorss species and environments, *h*_1_ = *h*_2_ = *s*_1_ = *s*_2_ (i.e., near the *global genotypic match* equilibria). See figure S7 for more information.

**Figure S10:**
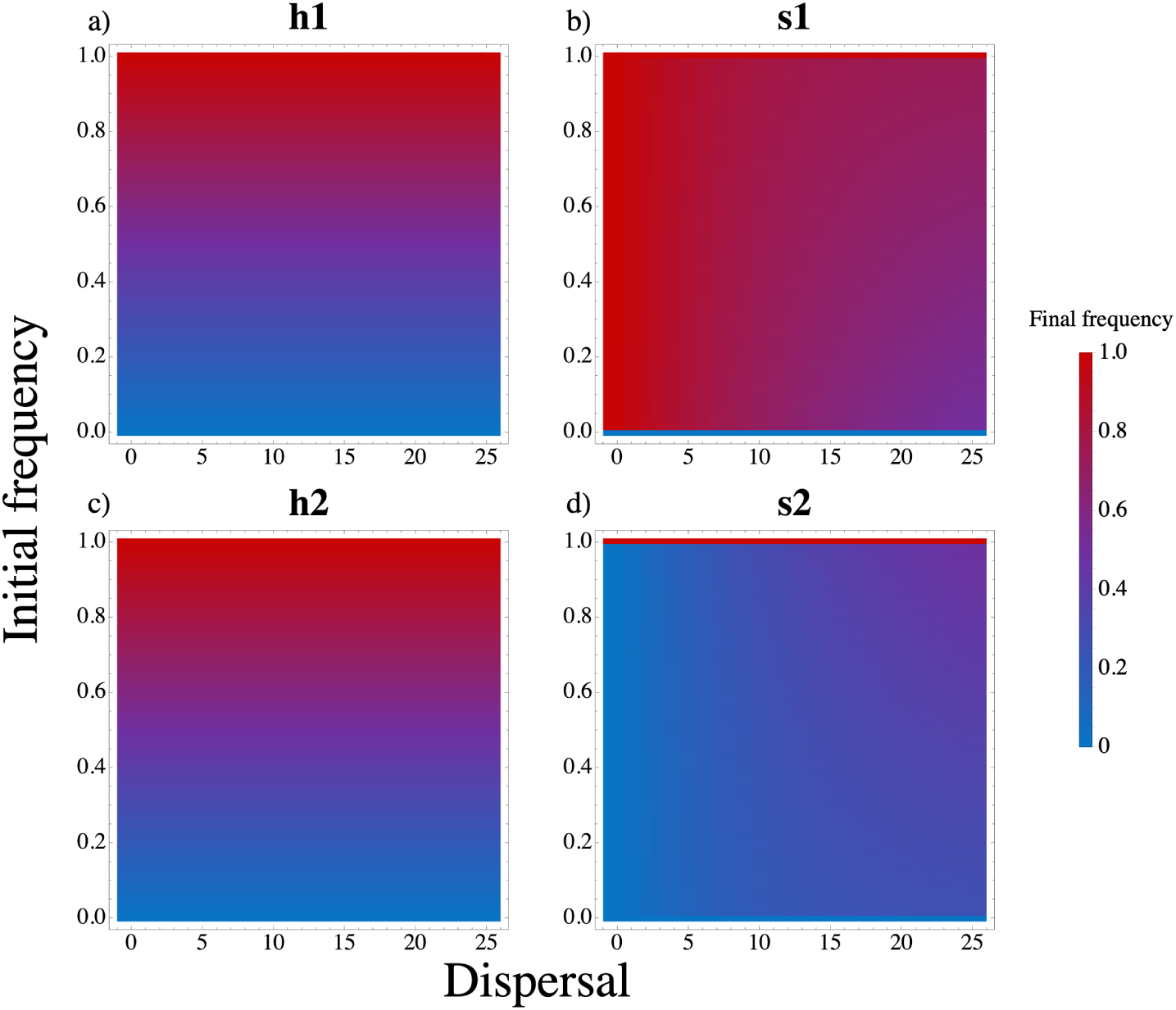
Results of simulations using parameters from Johnson et al. (2010), but averaged across each host and symbiont genotype. See figure S9 for more information.

**Figure S11:**
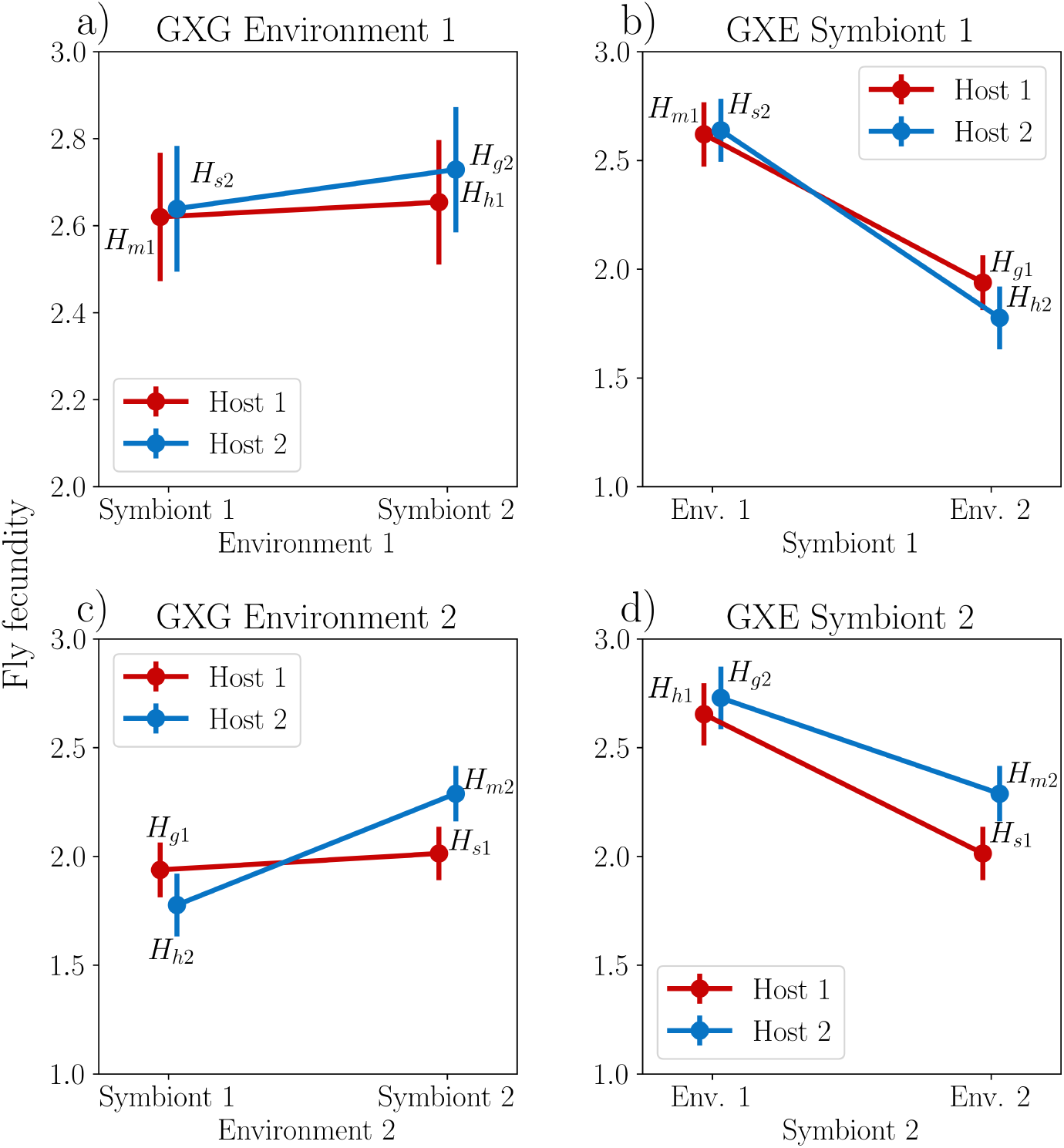
Empirical strengths of GXE and GXG in the *Drosophila melanogaster* microbiome (Henry et al., 2020). 1 denotes a control environment or genotype and 2 denotes a high sugar environment or genotype. Points are least squares means from Henry et al. (2020)’s statistical model and are labeled with their corresponding payoff (e.g. *H_h_*_1_ is host 1’s payoff from matching environment 1 and *H_h_*_2_ is host 2’s payoff from matching environment 2). Error bars represent the standard errors calculated in Henry et al. (2020).

**Figure S12:**
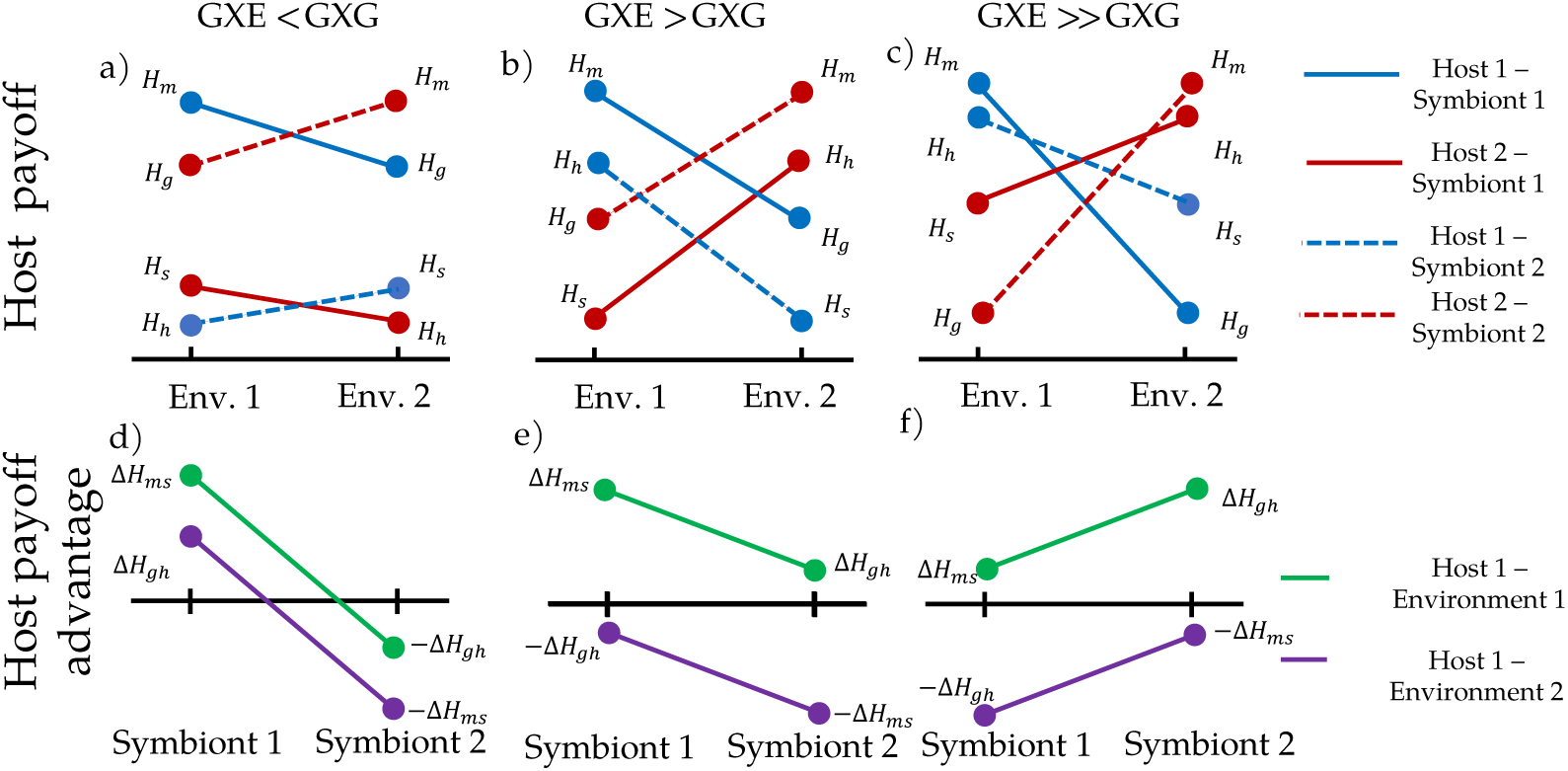
The relationship between the model’s payoffs and GXGXE interactions. The top row (a,b,c) shows payoffs between each environment for each host-symbiont combination. Line colors (blue or red) correspond to host genotype, while line styles (solid or dashed) correspond to symbiont gentoypes. The bottom row (d,e,f) shows the value of the payoff advantage for host 1 between symbiont partners. Green lines are environment 1 and purple lines are environment 2.

